# Expression partitioning of duplicate genes at single cell resolution in *Arabidopsis* roots

**DOI:** 10.1101/2020.08.20.260117

**Authors:** Jeremy E. Coate, Andrew D. Farmer, John Schiefelbein, Jeff J. Doyle

**Affiliations:** Department of Biology, Reed College, Portland, OR USA; National Center for Genome Resources, Santa Fe, NM USA; Department of Molecular, Cellular, and Developmental Biology, University of Michigan, Ann Arbor, MI USA; School of Integrative Plant Science, Plant Biology Section, Cornell University, Ithaca, NY USA

**Keywords:** gene duplication, single cell RNA-seq, cell type, cell state, polyploidy, expression subfunctionalization

## Abstract

Gene duplication is a key evolutionary phenomenon, prevalent in all organisms but particularly so in plants, where whole genome duplication (WGD; polyploidy) is a major force in genome evolution. Much effort has been expended in attempting to understand the evolution of duplicate genes, addressing such questions as why some paralogue pairs rapidly return to single copy status whereas, in other pairs, paralogues are retained and may (or may not) diverge in expression pattern or function. The effect of a gene—its site of expression and thus the initial locus of its function—occurs at the level of a cell comprising a single cell type at a given state of the cell’s development. Thus, it is critical to understand the expression of duplicated gene pairs at a cellular level of resolution. Using *Arabidopsis thaliana* root single cell transcriptomic data we identify 36 cell clusters, each representing a cell type at a particular developmental state, and analyze expression patterns of over 11,000 duplicate gene pairs produced by three cycles of polyploidy as well as by various types of single gene duplication mechanisms. We categorize paralogue pairs by their patterns of expression, identifying pairs showing strongly biased paralogue/homoeologue expression in different cell clusters. Notably, the precision of cell-level expression data permits the identification of pairs showing alternate bias, with each paralogue comprising 90% or greater of the pair’s expression in different cell clusters, consistent with subfunctionalization at the cell type or cell state level, and, in some cases, at the level of individual cells. We identify a set of over 7,000 genes whose expression in all 36 cell clusters suggests that the single copy ancestor of each was also expressed in all root cells. With this cell-level expression information we hypothesize that there have been major shifts in expression for the majority of duplicated genes, to different degrees depending, as expected, on gene function and duplication type, but also on the particular cell type and state.

## INTRODUCTION

According to Lynch and Trickovic (2020, p. 1861), “One of the last uncharted territories in evolutionary biology concerns the link with cell biology. Because all phenotypes ultimately derive from events at the cellular level, this connection is essential to building a mechanism-based theory of evolution.” As a candidate for building such a connection to cell biology, it would be difficult to identify a more important molecular evolutionary process than gene duplication, whose key role has been universally recognized since the classic paper of Ohno (1970), half a century ago. Gene duplication occurs at high frequency, estimated at 0.01 duplications per gene per million years in eukaryotic genomes, and large numbers of recently formed paralogues are found in typical animal, fungal, and plant genomes (Lynch and Conery, 2000; Lynch et al., 2001). Among eukaryotes, plant genomes, in particular, are characterized by massive levels of duplication, thanks to waves of whole genome duplication (WGD, polyploidy); recent estimates place the number of known plant WGD events at the genus level or above at over 250, and most plant lineages have experienced multiple cycles of polyploidy (Van de Peer et al., 2017; Leebens-Mack et al., 2019). Because they are the products of both small scale duplications (SSD) and WGD, plant gene families can be very large and complex (Panchy et al., 2016).

The fate of most paralogues, whether produced by SSD or WGD, is most likely to be pseudogenization and eventual loss, through mutations that inactivate redundant copies during the “fixation phase” of the gene’s life cycle (Innan and Kondrashov, 2010; Xie et al., 2019). Various mechanisms have been hypothesized that can preserve paralogue pairs by making both copies of the gene indispensable (Innan and Kondrashov 2010; Panchy et al., 2016; Qiao et al., 2019). These mechanisms can differ for SSD vs. WGD, even operating in different directions in the case of dosage effects (Freeling et al., 2009; Birchler and Veitia, 2014). Understanding why and how gene pairs are retained is complicated in part because in many cases competing hypotheses are difficult to distinguish from one another in terms of their predictions (Innan and Kondrashov, 2010). Obtaining empirical data for testing these hypotheses is not easy. Several of the models involve “function”, a term that can be difficult to define. Expression is often used as a proxy for function (e.g., Panchy et al., 2019), with biased expression of paralogues of a duplicated gene providing evidence for sub- or neofunctionalization.

Any phenotypic effects of gene duplication must necessarily manifest themselves at the level of single cells, and thus the processes that lead to the elimination or retention of paralogues operate initially at the cellular level, though ultimately the fate of a cellular innovation, like any mutation, is determined by the populational phenomena of genetic drift and selection (Lynch and Trickovic, 2020). Transcriptomic or proteomic studies, when conducted on organs or tissues, assay variation across multiple cell types and states (Efroni and Birnbaum, 2016; Libault et al., 2017). For understanding patterns of duplicate gene expression, this approach is problematic because it can miss cases of expression partitioning, by aggregating cells with expression fixed for different paralogues (Figure 1). Efforts have been made to circumvent this problem by analyzing expression at the level of single cell types, notably cell types for which large populations of pure cells can be obtained, such as seed fibers of cotton (*Gossypium*; e.g., Hovav et al., 2008a; Gallagher et al., 2020) or root hairs (e.g., Qiao and Libault 2013; Hossain et al., 2015), or through cell sorting in model systems such as *Arabidopsis* (Birnbaum et al., 2003). However, even when samples comprising only a single cell type can be analyzed, these can contain multiple cell developmental states, and “the” transcriptome or proteome thus obtained will again be the sum of more than one potentially differentiated sets of transcripts or proteins. Additionally, by focusing on a single cell type, such studies are blind to any expression partitioning between paralogues that might have occurred across cell types.

**Figure 1.**
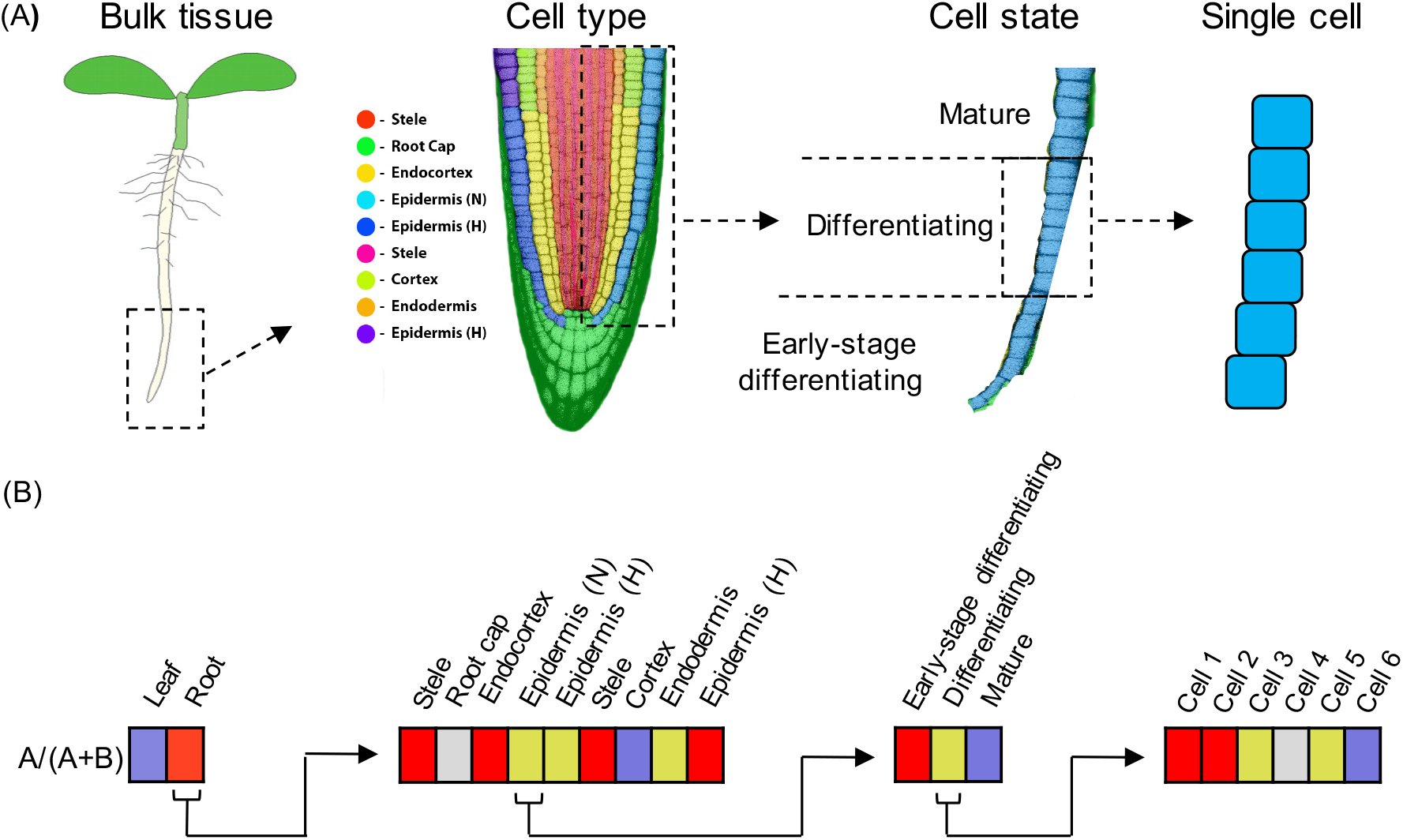
**(A)** Expression subfunctionalization is the partitioning of ancestral expression profiles between paralogues. Partitioning can occur at the level of tissue types (e.g., leaves vs. roots), but also at increasingly finer scales, including at the level of cell type (in roots, roughly corresponding to the nine cell clusters identified by Ryu et al., 2019), or cell state (e.g., developmental gradients within a single cell type such as non-hair epidermal cells), and even among single cells. **(B)** Examples of how expression partitioning between paralogues (designated “A” and “B”) can occur at different levels. Expression bias is indicated by shading (red = expression biased towards paralogue A [specifically, A/(A+B)>0.9], blue = expression biased towards paralogue B [A/(A+B)<0.1], yellow = unbiased expression, grey = neither paralogue expressed). Left, In this example, paralogue A expression predominates in roots, whereas paralogue B expression predominates in leaves. Middle left, Within root tissue, paralogues A and B exhibit partitioning by cell type. Middle right, within the non-hair epidermal cell type, paralogues A and B exhibit partitioning by cell state (i.e. developmental stage). Right, within the “differentiating” stage of non-hair epidermal cell type, A and B paralogue expression is partitioned among individual cells (paralogue A predominating in cells 1-2, and paralogue B expression predominating in cell 6).

Single cell methods that already have revolutionized biology continue to develop and promise ever more powerful and precise data (Lähnemann et al., 2020). In plants, several groups recently published single cell transcriptomic studies of *Arabidopsis* roots (Jean-Baptiste et al., 2019; Ryu et al., 2019; Shulse et al., 2019; Denyer et al., 2019; Zhang et al., 2019) that not only identified the expected cell types, including cell types represented by small numbers of cells that would be missed in conventional transcriptomic studies, but also revealed cells with distinctive transcriptomes not readily assigned to known cell types. These data represent a resource for understanding expression of duplicate genes at an unprecedented level of detail.

Here we elucidate patterns of expression in 36 root cell clusters representing root cell types and their developmental states for over 11,000 *Arabidopsis* paralogue pairs produced by various types of duplication (Wang et al., 2013). We identify a set of pairs that show strong bias in paralogue expression in root cell types and states, including some for which bias shifts from one paralogue to the other among cell types. We also identify a set of over 7,000 genes expressed in all root cell types, which we infer to be essential for root cell function, and explore the degree to which paralogues, particularly those created by the alpha WGD that took place around 30-40 million years ago (MYA) in the ancestor of Brassicaceae (Edger et al., 2018), either share this ubiquitous expression pattern or exhibit shifts in expression from the presumed ancestral state. We find that many pairs of alpha WGD homoeologues show evidence of expression shifts, but not to the degree observed for SSD pairs. Interestingly, different cell types vary considerably in the degree to which they express both paralogues of these gene pairs.

## METHODS

### Single cell datasets

Illumina sequence data for NCBI SRA experiments SRX5074330-SRX5074332 corresponding to scRNA-seq data from three wild-type replicates of *Arabidopsis* roots from Ryu et al. (2019) were processed with the 10X Genomics Cell Ranger v3.1.0 count pipeline, run independently on the data for each replicate to produce unique molecular identifier (UMI) raw counts matrices against the TAIR10 genome using Araport11 annotations (Cheng et al., 2017). Custom scripts (available from https://github.com/adf-ncgr/singlecell_paralogue_expression_scripts) were used to produce per-cluster UMI counts for each gene, summing the contributions from all cells assigned to the 9 “superclusters” presented in Ryu et al. (2019) and separately for 36 root cell clusters (“RCCs”) derived from those 9 initial superclusters using the Seurat software package (Butler et al., 2018; FindClusters function) with default parameters (perplexity=30, random seed=1) and a resolution of 3.5. These per-supercluster and per-RCC gene UMI count matrices formed the basis of subsequent analysis of expression bias between duplicated gene pairs. Specific cell types and differentiation states were assigned to each of the 36 RCCs from the 9 initial superclusters (Supplemental Table 1) using previously defined marker genes (Ryu et al., 2019).

To filter out spurious expression signals (resulting, for example, from doublets or from cell-free RNA), we required UMI from two or more cells in a given cluster for a gene to be considered expressed in the context represented by the cluster. In some cases, we also analyzed the data using the minimally restrictive expression threshold of ≥1 UMI from ≥1 cell to assess how strongly additional filtering affects the results.

The cells in the Ryu et al. (2019) study from which the data were taken were derived from protoplasted root tissues, and would thus be subject to some level of protoplasting-induced changes in gene expression relative to untreated tissues (Birnbaum et al., 2003). We considered that altered responses to the protoplasting treatment were within the scope of what could be considered divergence, and retained duplicate pairs involving such genes in the subsequent analyses.

### Sequence Read Archive (SRA) datasets

A total of 214 RNA-seq datasets for *Arabidopsis thaliana* used in Panchy et al. (2019) were obtained from NCBI SRA (Leinonen et al., 2011) and aligned to the TAIR 10 genome using hisat2 v2.1.0 (Kim et al., 2019) against an index built with splice sites and exons derived from Araport11 annotations. TPM values derived by stringtie v2.0.6 (Kovaka et al. 2019) were then converted to raw counts for the Araport 11 genes (Supplemental Table 2). These data were used to assess if genes not expressed in the Ryu et al. (2019) scRNA-seq data were expressed in other tissues and/or conditions. Of the 214 SRA libraries, 37 were generated from leaf tissue only (832 million reads total) and 31 were from root tissue only (290 million reads total). Counts were summed across all samples in each class (leaf vs. root) to provide high-coverage bulked data sets to compare paralogue expression at the level of contrasting tissue types, in the same manner as for the clustered single-cell data, as described below.

### Biased expression of paralogues

For gene pairs identified by Wang et al. (2013), we determined whether the paralogues showed biased expression (analogous to “biased homoeologue expression” of Grover et al., 2012) in the context of each cluster, using a UMI cutoff of 9:1. We chose 9:1 as a stringent threshold for biased expression because our primary interest was in identifying cases of extreme imbalance in paralogue expression, consistent with expression subfunctionalization. The two paralogues of the gene pair were then designated “single-cell A” (scA) and “single-cell B” (scB), with scA being the dominant paralogue showing higher expression in the greatest number of clusters.

In order to exclude from consideration gene pairs whose apparent bias may be insignificant relative to random sampling deviations, we considered the counts characterizing each gene pair as representing the outcome of a Bernoulli trial, where the abundance of reads relative to its partner determines the probability of each outcome (i.e., sampling a particular partner when a read is chosen from one of the pair). In order to determine confidence in the estimate of the probability given a specific number of trials (i.e., the summed read count for the pair), the Wilson (1927) score interval estimation was used to provide a 95% confidence interval around the expression bias value estimated from the read counts. Considered from the perspective of the dominant gene in a putatively biased pair, the lower bound on the confidence interval can be used as the minimum level of bias at the chosen confidence level, and if this minimum level of bias falls below the 9:1 threshold ratio for considering a gene pair to exhibit bias, it was removed from consideration when calculating the fixation and balance indices described below.

For each paralogue pair we calculated two indices to describe its pattern of expression across cell clusters:

- The Expression Fixation index (F_ex_) measures the degree of bias in the expression of paralogues of a given pair across the cell clusters. F_ex_ = N_fix_/(number of cell clusters expressing at least one paralogue above a cutoff threshold), where N_fix_ is the number of clusters for which one paralogue (either one) is “fixed” (is preferentially expressed at or above the 9:1 threshold).
- The Balance Index (B_fix_) is calculated for any paralogue pair for which at least one cluster is fixed for a paralogue, and measures the degree to which one paralogue dominates the expression across the clusters. B_fix_ = 2*(number of clusters fixed for scB)/(number of clusters fixed for either paralogue). B_fix_ runs from 0-1; cases with no fixation of either paralogue (paralogue pairs with F_ex_ = 0) have no B_fix_ score (N/A). Examples are given in Figure 2.

**Figure 2.**
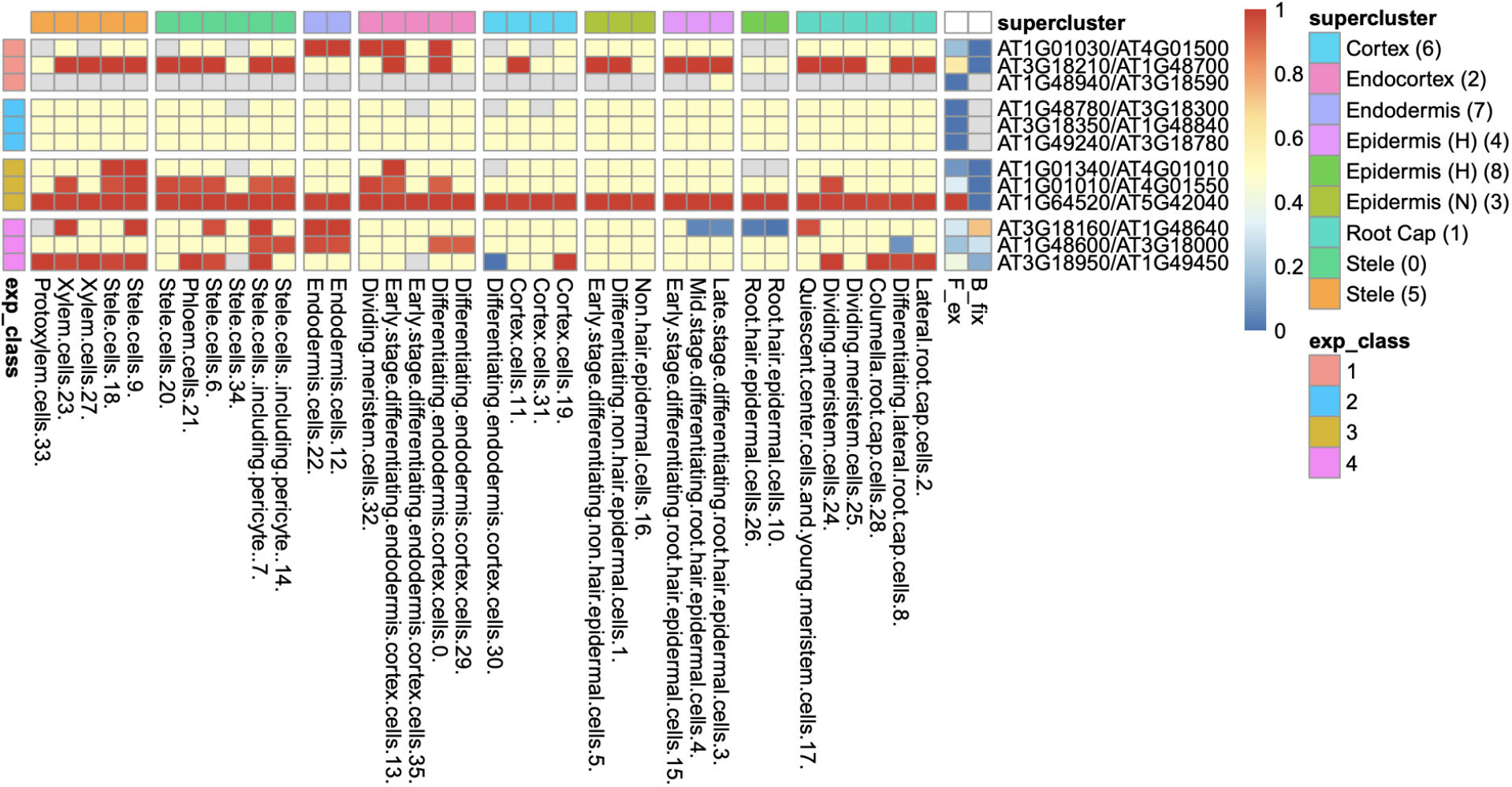
Heatmap of 36 root cell clusters (RCCs) from nine Ryu et al. (2019) superclusters, showing examples of selected gene pairs illustrating F_ex_ and B_fix_ (last two columns before gene pair designations) and the four expression Classes (defined in Table 1). Cell colors: For RCCs, red cells have dominant paralogue scA fixed (expressed above the 9:1 threshold at 95% confidence level, or, in the case of Class 1, expressed at 1:0); blue cells have paralogue scB fixed; yellow cells have at least one of the paralogues expressed, with neither fixed; gray cells have no expression of either paralogue. For F_ex_ and B_fix_ cell values for each index run from 0-1 and are colored continuously from blue to red. Examples of calculation of F_ex_ and B_fix_ for Classes 1-4: Rows 1-3 are Class 1 pairs, meaning that only one paralogue is expressed in any RCC; row 1 (AT1G01030/AT4G01500) has expression in 28/36 RCCs, with paralogue scA fixed in 5 and no fixation of scB, so F_ex_ = 5/28 = 0.18, and B_fix_ = 2(0)/5 = 0; Row 2 has expression in all 36 RCCs, scA fixed in 22 and none fixed for scB, so F_ex_ = 22/36 = 0.61, and B_fix_ = 2(0)/22 = 0. Class 2 pairs (rows 3-6) have both paralogues expressed in at least one RCC, but no fixation of either paralogue in any cluster, so F_ex_ = 0 and B_fix_ is not calculable (gray). Class 3 genes (rows 7-9) have both paralogues expressed in one or more RCCs and have scA fixed in one or more RCCs, with scB never fixed (so B_fix_ = 0 for all three pairs); the row 7 pair has expression in 32 RCCs and fixation of scA in 3, F_ex_ = 3/32 = 0.09; in row 8, F_ex_ = 12/36 = 0.33; in row 9, F_ex_ = 36/36 = 1.0. Class 4 pairs have both paralogues fixed in different RCCs, so F_ex_ > 0, and B_fix_ > 0; for example, in row 10, the pair has at least one paralogue expressed in 35 RCCs, with 7 RCCs fixed for scA (red) and 4 RCCs fixed for scB (blue), so F_ex_ = 11/35 = 0.31, and B_fix_ = 2(4)/11 = 0.73.

**Table 1.**
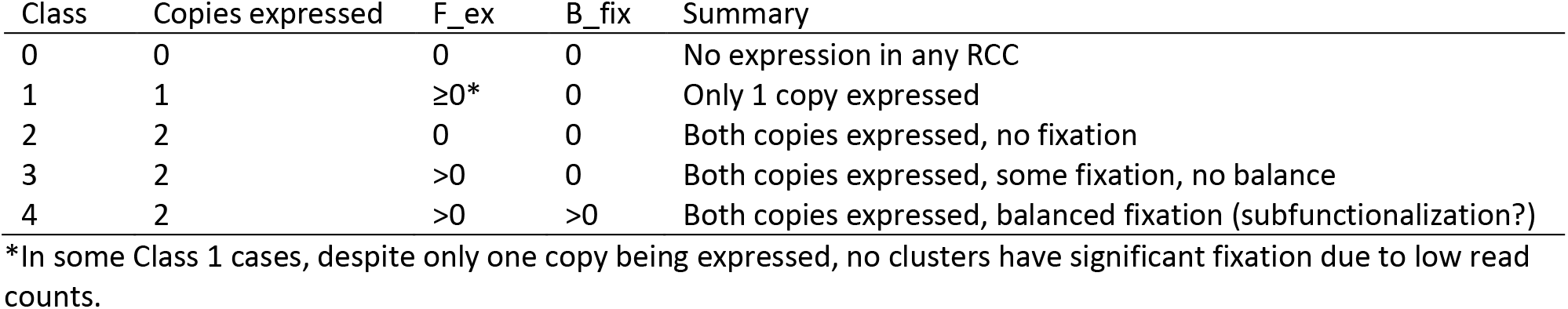
Expression classes of duplicate gene pairs.

| Class | Copies expressed | $F_{\text{ex}}$ | $B_{\text{fix}}$ | Summary |
| --- | --- | --- | --- | --- |
| 0 | 0 | 0 | 0 | No expression in any RCC |
| 1 | 1 | $\geq 0^*$ | 0 | Only 1 copy expressed |
| 2 | 2 | 0 | 0 | Both copies expressed, no fixation |
| 3 | 2 | $> 0$ | 0 | Both copies expressed, some fixation, no balance |
| 4 | 2 | $> 0$ | $> 0$ | Both copies expressed, balanced fixation (subfunctionalization?) |

\*In some Class 1 cases, despite only one copy being expressed, no clusters have significant fixation due to low read counts.

### Classification of paralogues by expression pattern

We classified paralogue pairs into five classes based on their expression patterns across the 36 RCCs (Table 1; Figure 2). Genes were also assigned to equivalent classes at the nine cluster level (e.g., Class 1 if only one of the two copies was expressed at the 9-supercluster level).

### Fixation similarities and differences across clusters

Heatmaps were produced on the results of calculating biased expression for each gene pair across all clusters, by using the R package “Pretty Heatmaps” (pheatmap v1.0.12). The annotations option of pheatmap was used to denote the expression class (Table 1).

### Single cell measurements of paralogue usage

In order to look at possible bias between duplicated genes at the level of single cells, we again applied the Wilson (1927) test on the UMI counts for the duplicated genes at the level of individual cells. In this case, due to the low counts obtained and in order not to confute subsequent tests we extended the states recorded by the test to distinguish not only between biased and unbiased but also cases where the counts were too low to determine whether a particular gene pair fell in one class or another in a given cell. Cells in the latter case were excluded from subsequent analyses. For the remainder, gene pairs could be tallied with respect to the number of cells in a given cluster that showed bias for one or the other of the genes, were unbiased with respect to the expression of the genes, or showed no expression of either gene (i.e., had a UMI count of zero for both). The number of cells falling into each category were then used to test for significant overlap of the lists of cells expressing each the two genes, using Fisher’s exact test in the manner of the GeneOverlap Bioconductor package (Shen and Sinai, 2019), while additionally testing for significant non-overlap (i.e. a tendency for cells expressing one paralogue to not express the other paralogue) by utilizing the alternate tail of the hypergeometric distribution.

### Ka/Ks value determination for paralogues

In order to assess whether different classes of paralogues based on patterns of expression showed significant differences in terms of protein coding divergence, we used the implementation of the Nei-Gojobori algorithm (Nei and Gojobori, 1986) provided by the Bio::Align::DNAStatistics module of BioPerl (Stajich et al., 2002) through wrapper scripts available in the MCScanX software distribution (Wang et al., 2012) but run across all the *Arabidopsis* gene duplicate pairs classified by Wang et al. (2013) including non-syntenic SSD pairs.

## RESULTS

### Arabidopsis root cell clusters each express over 35% of genes in the genome

We studied both the 9 root cell supercluster transcriptome data of Ryu et al. (2019) and transcriptomes of the 36 root cell clusters (RCCs) derived from those superclusters. These two datasets are strongly nested, with the 9 superclusters broadly corresponding to cell types and the 36 RCC data including developmental states of those types (Figure 3, Table 1). We primarily report results from the 36 RCCs here (9 supercluster data and 36 RCC counts data are available in Supplemental Table 3).

**Figure 3.**
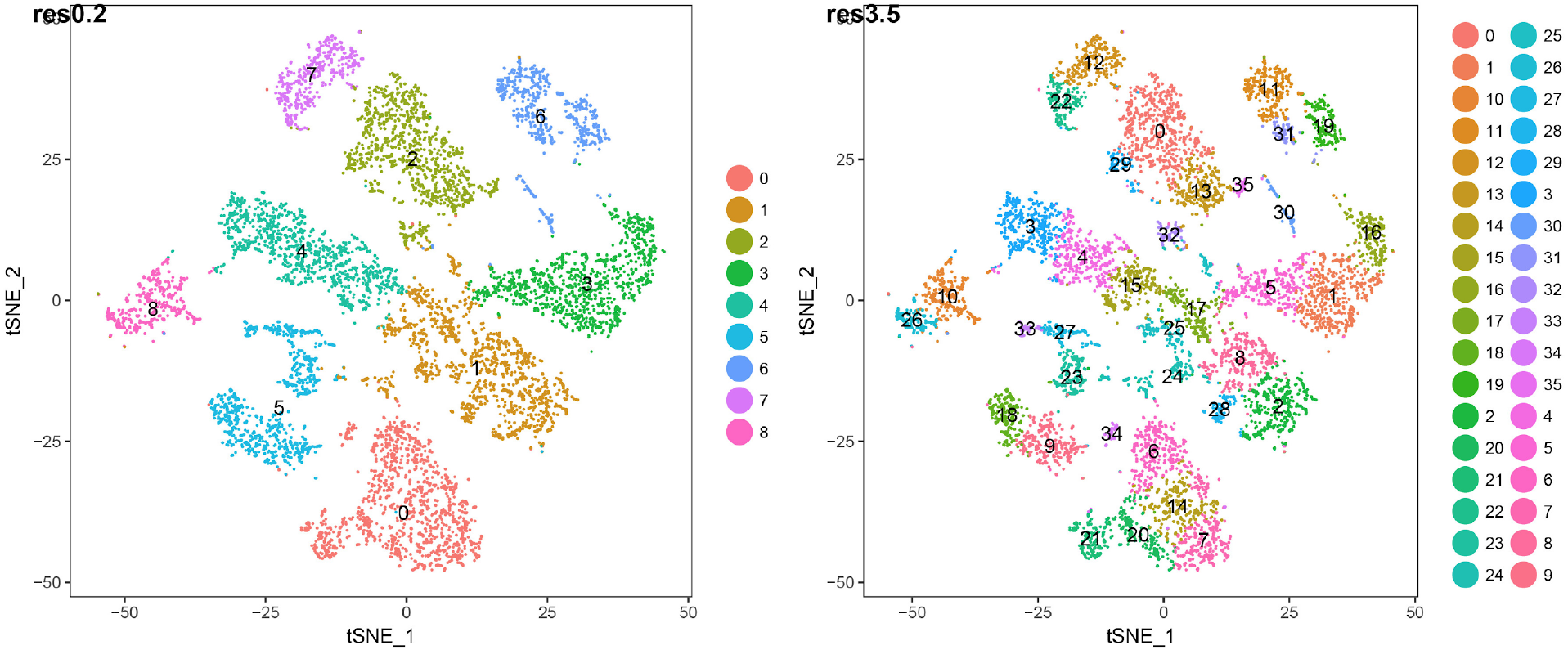
t-SNE plots of 7522 wild-type *Arabidopsis* root single-cell transcriptomes clustered at two different resolutions. The lower resolution generates 9 major clusters (“superclusters”; roughly representing “cell types”) and the higher resolution generates 36 root cell clusters (RCCs; roughly representing “cell states”). The specific cell type/state assignments for each cluster is provided in Figure 2. Additional details are provided in Supplemental Table 1.

We explored two different thresholds for determining whether a gene was expressed in an RCC: at least 1 UMI per cluster vs. the more stringent cutoff of expression in two or more cells of a given RCC. These thresholds were chosen to accommodate different technical issues with single-cell RNA-seq droplet-based methods (Leucken and Theis, 2019). On the one hand, technical dropout (Bhargava et al., 2014) leads to reduced capture for low abundance transcripts, suggesting a relaxed threshold for counts relative to bulk RNA-seq; conversely, the possibility of capturing multiple cells of differing types in a single droplet (“doublets”) suggests a need to guard against false positives generated by this phenomenon. We found that counts derived from these two thresholds were strongly correlated (R^2^ = 0.997) but 7-18% lower for the two cell cutoff than for the 1 UMI cutoff (Supplemental Figure 1); we report numbers using the more stringent cutoff throughout.

After excluding loci annotated as “novel transcribed regions”, “pseudogenes”, “transposable element genes”, and organelle-encoded genes, there are 32,548 genes in the most recent annotation of the *Arabidopsis* genome (Araport11: Cheng et al., 2017). Of these, 22,669 (70%) were expressed in at least one of the 36 RCCs, with each RCC expressing 35-58% of these genes (Figure 4). These percentages are comparable to *Arabidopsis* pollen cell stages (32-51% of microarray features; Honys and Twell, 2004), and are somewhat lower than those for cotton fiber cells, which transcribe from 75-94% of the genome’s genes depending on developmental stage (Hovav et al., 2008b). Differences in the number of genes expressed per cell cluster were in many cases statistically significant, but these differences were largely driven by differences in cell count per RCC (Supplemental Figure 2). Approximately 35% of root-expressed genes (^~^22% of all *Arabidopsis* genes) were expressed in all 36 RCCs (referred to as “RCC-ubiquitous” or “RCC-u”); 1,059 genes (4.7% of root-expressed genes) were uniquely expressed in only one RCC (Figure 4).

**Figure 4.**
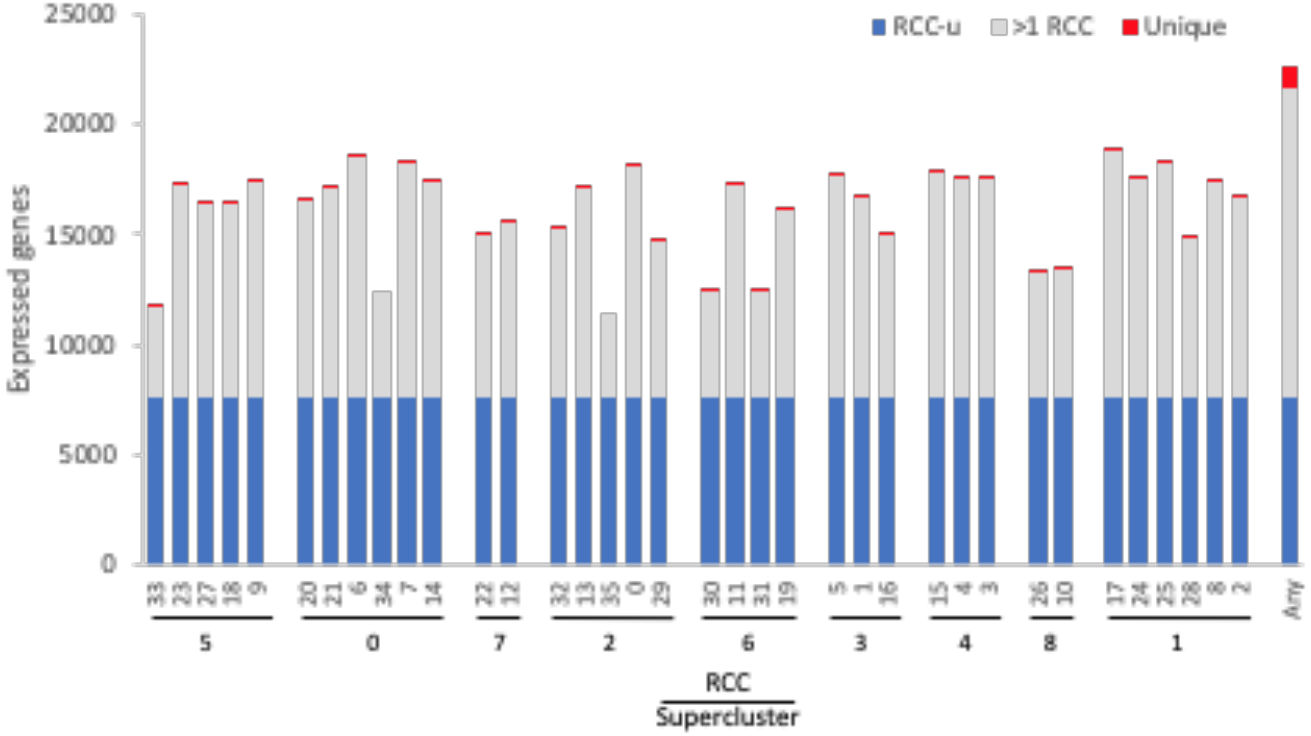
Number of genes expressed per cluster. The 36 RCCs are grouped by Ryu et al. (2019) supercluster (roughly representing cell types and developmental states; Figure 1), and ordered roughly from the interior (stele) to the exterior (epidermis) and tip (root cap) of the root. RCC-ubiquitous (RCC-u) is the subset of genes expressed in all 36 RCCs, “>1 RCC” is the subset expressed in 2-35 RCCs, and “Unique” is the subset expressed in only one RCC. “Any” is the union of all genes expressed in at least one of the 36 clusters.

### Many gene pairs show biased paralogue expression in root cell clusters, and different duplication types show different expression patterns

Wang et al. (2013) identified 11,470 gene pairs in the Arabidopsis genome, and classified them as being duplicated either by WGD or by SSD. They further subdivided the WGD class into products of the alpha (most recent, around 31.8-42.8 MYA; Edger et al., 2018), beta (85-92.2 MYA; Edger et al., 2018), and gamma events (115-120 MYA; Jiao et al., 2012), and the SSD class into tandem duplicates, proximal duplicates, and two subclasses of transposed duplicates (younger than 16 MY vs. older).

The expression breadth of pairs (number of clusters in which a pair is expressed) differed by duplication type (Supplemental Figure 3). For all three WGD types, around 44-48% of paralogue pairs had one or both paralogues expressed in all 36 RCCs (“RCC-u pairs”), and 30-38% of genes were RCC-u. Only 4-6% of pairs had neither paralogue expressed in any RCC (i.e., Class 0 pairs; Table 1, Figure 5, Supplemental Figure 3), comprising 8-12% of all genes. In contrast, the reverse was true for tandem and proximal SSD types, with 25-30% of pairs and around 35-40% of their individual genes not expressed in any RCC and fewer than 20% of pairs and 12% of their genes expressed in all 36 RCCs. Transposed duplicates were intermediate between these two groups, with older pairs behaving more like WGD types (44% of pairs and 30% of individual genes expressed in all RCCs; 2% of pairs and 12% of genes not expressed in any RCC) and younger transposed pairs behaving more like the other SSD types (28% of all pairs and 17% of genes expressed in all 36 RCCs; 18% of pairs and 33% of genes not expressed in any RCC). As noted by Wang et al. (2013), on average, based on K_s_, proximal and tandem duplicates are the youngest of the duplicated pairs, and the lower percentages of generally younger duplicate pairs expressed in root cell clusters may be due to the difficulty in mapping short reads unambiguously to paralogues with very similar sequences, a problem noted by others (Wang et al., 2013; Lan and Pritchard, 2016; Defoort et al., 2019).

**Figure 5.**
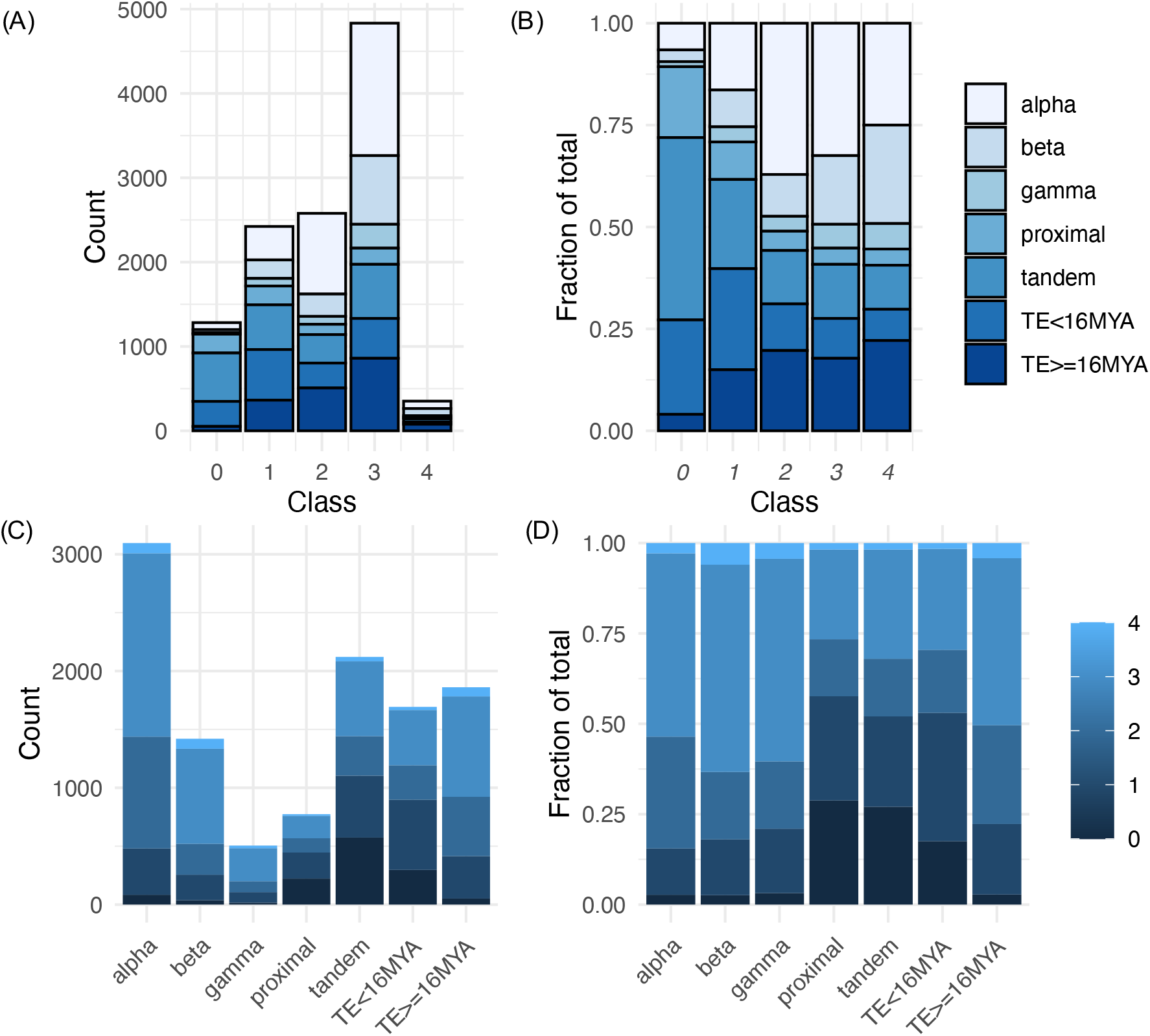
Distribution of expression classes and duplication types for gene pairs. **(A)** Counts of gene pairs by expression class, broken down by duplication mechanism. **(B)** Fraction of gene pairs in each expression class produced by each duplication mechanism. **(C)** Counts of gene pairs by duplication mechanism, broken down by expression class. **(D)** Fraction of gene pairs from each duplication mechanism assigned to each expression class.

We divided these 11,470 paralogue pairs into five RCC expression classes according to whether neither paralogue (Class 0), only one paralogue (Class 1), or both paralogues (Classes 2, 3 and 4) were expressed in at least one cluster; Classes 2, 3 and 4 were distinguished by patterns of biased paralogue expression estimated at a 9:1 ratio (Figure 2). Overall, 10,187 of these gene pairs (88.8% of total pairs) belonged to Classes 1-4, having one or both paralogues expressed in at least one RCC (Supplemental Table 4). This percentage is similar to but significantly lower than the expectation for drawing at least one member of a gene pair from the 70% of *Arabidopsis* genes expressed in root cell clusters (90.8%; *χ*^2^ = 24.3, p < 0.001). Notably, however, 67.7% (7,751 pairs) had both copies expressed in root cell clusters, significantly higher than the random expectation of 48.5% (*χ*^2^ = 856.9, p < 0.001). There was a clear distinction between WGD and local SSD (proximal and tandem) pairs with regard to these percentages. WGD pairs were significantly more likely to express at least one copy (≥96.8%; p < 0 0.01) and to express both copies (≥79.0%%; p < 0.01) than expected by chance, whereas local SSD pairs were significantly less likely to express at least one copy (≤72.9%; p < 0.01) and less than or similarly likely to express both copies (≤47.9%). Older duplicates created by transposable elements (TEs) exhibited a similar pattern to WGD duplicates and younger TEs exhibited a similar pattern to local SSD duplicates (Figure 5, Supplemental Table 4).

Class 1 gene pairs have the same paralogue exclusively expressed in all root cell clusters in which either paralogue is expressed (balance index [B_fix_] = 0) (Figure 2; Table 1). The same dichotomy between duplication types observed for Class 0 gene pairs was also observed for Class 1: Only around 10-20% of WGD and older transposed duplicate pairs belonged to this class, vs. 25-35% of tandem, proximal, and younger transposed SSD classes (Figure 5, Supplemental Table 4).

In a Class 1 pair, only the dominant paralogue (scA) is expressed; scB shows no expression in any RCC, and could either be a pseudogene, or could be a functional gene expressed in other contexts than the roots studied here. We thus looked at expression for genes of this class in 214 RNA-seq experiments obtained from the SRA database (Leinonen et al., 2011) (Supplemental Table 2). We found that in 93% of Class 1 gene pairs, the scB paralogue was expressed in at least one SRA dataset, and 91% were expressed in non-root SRA libraries (requiring at least 5 reads to be considered expressed) (Supplemental Table 5). These results indicate that the majority of Class 1 genes (90-93%) represent cases where expression has been partitioned between paralogues since their divergence, with scA being the only copy now expressed in roots grown under the conditions used by Ryu et al. (2019). Though this suggests that many of these scB paralogues are functional, some could be pseudogenes, because around a third of plant pseudogenes are expressed (Xie et al., 2019). The 7-10% of Class 1 pairs for which there is no evidence of scB expression in any SRA libraries have elevated Ka/Ks (mean = 1.41, median = 0.92) relative to pairs for which scB is expressed (mean = 0.65, median = 0.39) (t = −7.3, df = 148, p < 0.001), indicative of relaxation of selection, consistent with pseudogenization but also with positive selection.

In pairs comprising Classes 2-4, both paralogues are expressed in at least one cluster, with Classes 3 and 4 distinguished from Class 2 by biased paralogue expression defined at the stringent threshold of 9:1, and Class 3 and 4 pairs differentiated by whether only scA showed bias (Class 3) or both scA and scB showed biased expression in different RCCs (Class 4; Table 1). Whereas the majority of Class 0 and Class 1 pairs are from proximal, tandem or young transposed duplicates, the majority of pairs in Classes 2-4 were produced by WGD or older transpositions (Figure 5). This pattern was most pronounced for Class 3, in which both copies are expressed but scA, and only scA, is expressed in at least one RCC above the bias cutoff (Figure 5). Class 4 pairs (pairs in which both copies are expressed, with scA and scB expressed above the bias cutoff in different clusters) comprised by far the smallest number of pairs (only 1.5-5.6% of pairs in duplication types), but showed the same pattern as other pair classes, with WGD and older transposed pairs having a larger percentage representation than the other SSD types.

Expression classes aggregate data across individual RCCs to summarize expression patterns, but do not provide information about paralogue behaviour within individual RCCs. Consequently, we also looked at levels of bias within RCCs. Different classes of duplication also showed very different percentages of pairs exhibiting bias in individual RCCs, with proximal, tandem, and younger transposed classes all showing greater levels of bias than WGD and older transposed pairs (Figure 6). Homoeologues from alpha WGD pairs are the least likely to show expression bias. The fraction of biased pairs also varied by RCC, with root cap clusters generally exhibiting the least bias across all types of duplicates.

**Figure 6.**
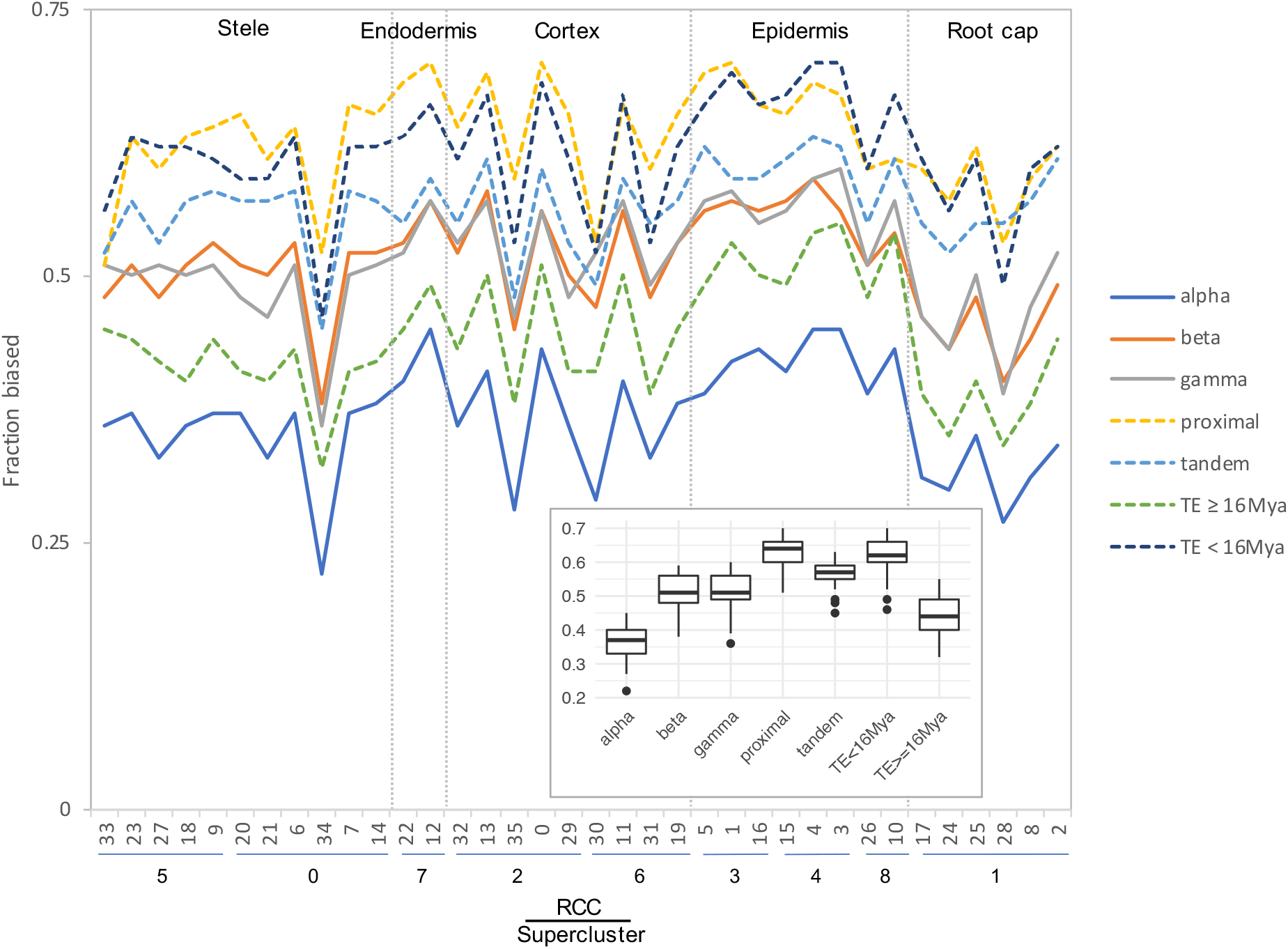
Fraction of gene pairs exhibiting paralogue expression bias by RCC. RCCs are arranged by supercluster and ordered roughly from the interior (stele) to the exterior (epidermis) and tip (root cap) of the root. Each line indicates the fraction of gene pairs from the specified duplication mechanism that exhibit expression bias per RCC. For ease of comparison, whole genome duplications are shown with solid lines, and small scale duplications are shown with dashed lines. Individual gene pair by RCC combinations that lacked sufficient read depth to detect bias were excluded from the analysis. Inset, box plot summarizing the distribution of bias fractions by duplication type.

### Genes showing “alternate fixation”

Class 4 pairs show extreme reciprocal expression biases across clusters, and are the most likely cases of shifts in function following duplication. To determine how much greater resolution to detection functional differentiation was provided by cell-level data, we determined the number of Class 4 genes: 1) between bulk tissues (root vs. leaf SRA libraries); 2) across the 9 superclusters, roughly representing cell types; 3) across the 36 RCCs, further subdividing cell types into cell states; and 4) between individual cells of the 36 RCCs (Figure 7).

**Figure 7.**
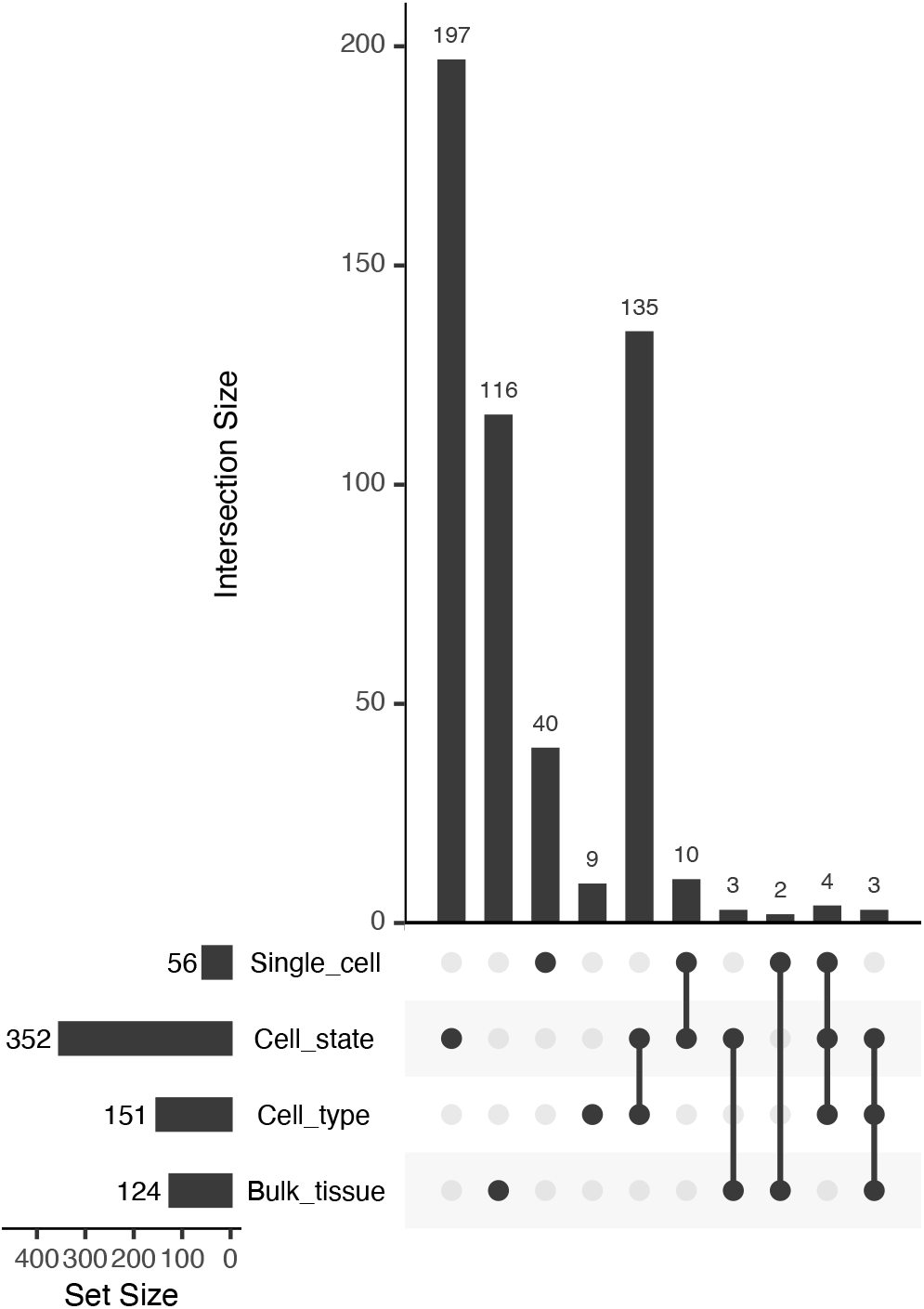
UpSet plot of intersections among Class 4 genes identified at different levels of organization.

The number of cases of paralogue expression bias identified using the single cell data is three times greater than those identified in the bulk tissue comparison (403 vs. 124; Figure 7). Most of these cases (352) were identified at the level of the 36 RCCs, suggesting that in root tissue paralogue expression bias occurs more frequently among cell states within a cell type than among cell types, with the caveat that the 9 superclusters may include more than one cell type.

Within the 9 superclusters, only 56 gene pairs exhibited paralogue expression bias at the level of single cells (an over-representation of cells exhibiting significant bias favoring one copy in some cells, and the other copy in other cells, with few or no cells co-expressing both equally). This is likely an undercount of the true number, however, due to the low read count per cell. Of the 87.5 million possible cell x gene pair comparisons (7,519 cells × 11,631 gene pairs), 19.2 million had non-zero read counts, but of these, 17.5 million had counts that were nonetheless too low to detect bias at our threshold of 9:1. In contrast to 56 significantly non-overlapping gene pairs, 1,363 gene pairs exhibited significant overlap at the level of single cells (cells expressing one copy were significantly more likely to express the other copy as well). Alpha WGD duplicates were the most likely to exhibit significant overlap at the level of single cells, whereas gamma WGD duplicates were the most likely to exhibit significant non-overlap (Table 2).

**Table 2.** Counts of overlap and non-overlap of paralogue expression in individual cells.

| Duplication type | Overlap | % of total | Non-overlap | % of total |
| --- | --- | --- | --- | --- |
| alpha | 593 | 18.64% | 13 | 0.41% |
| beta | 170 | 11.72% | 5 | 0.34% |
| gamma | 64 | 12.28% | 4 | 0.77% |
| proximal | 39 | 4.97% | 2 | 0.26% |
| tandem | 149 | 7.00% | 12 | 0.56% |
| TE<16 MYA | 103 | 6.06% | 7 | 0.41% |
| TE≥16 MYA | 245 | 13.16% | 13 | 0.70% |

Notably, Class 4 genes at all four levels of organization (bulk tissue, cell types, cell states, and single cells) are enriched for extracellular functions (e.g., extracellular region, apoplast, cell-cell junction), and each is also enriched for some aspect of the cell periphery (e.g., cell wall, plasma membrane). Thus, balanced fixation of expression, suggestive of partitioning of function, appears to occur preferentially among paralogues functioning at the cell surface. Beyond this commonality, however, GO enrichment analysis suggests that paralogues exhibiting balanced fixation at the bulk tissue level differ functionally from those exhibiting balanced fixation at finer levels of resolution. At the bulk tissue level, Class 4 genes are preferentially involved in lipid metabolism and vesicle trafficking (exocyst), whereas at the supercluster, RCC and single cell levels, Class 4 genes are preferentially involved in cell wall modification (e.g., cell wall organization or biogenesis, hemicellulose metabolic process) and response to stress (e.g., response to oxidative stress, response to toxic substance).

### Concerted divergence of paralogous genes

Overall, WGD gene pairs were more likely to exhibit balanced expression bias (alternate fixation across the 36 RCCs; Class 4) than were SSDs (X^2^ = 22.1, p < 0.001). In total, 195 out of 5,018 WGD pairs (3.9%) were assigned to Class 4, compared to 157 out of 6,449 SSD pairs (2.4%).

It has been proposed that WGD facilitates functional differentiation by simultaneously duplicating entire gene networks, thereby providing the raw material for concerted whole pathway evolution (“concerted divergence”: Blanc and Wolfe, 2004). We looked for evidence of concerted divergence, focusing on the genes duplicated by the alpha WGD. Out of a total of 3,096 alpha gene pairs, 176 genes in 88 pairs exhibited alternate fixation across the 36 RCCs (Class 4), indicative of expression subfunctionalization.

For each of these 88 Class 4 alpha pairs we calculated an expression ratio (scA/total) for each of the 36 RCCs. Then, for each alpha pair, we calculated correlation coefficients with every other alpha pair based on these 36 expression ratios. To the extent that two alpha pairs have diverged in concert, we would expect their expression ratios to be either positively correlated (r ≫ 0) if scA from both pairs are coevolving, or negatively correlated (r ≪ 0) if scA from one pair is coevolving with scB from the other pair. For alpha pairs that are diverging independently of each other, we expect to see no correlation (r ≈ 0). By this approach, 33 of the 88 pairs formed a distinct set of significantly correlated expression ratios (p < 0.05; Figure 8A), suggesting concerted divergence into two separate networks. Several additional, smaller clusters were evident as well.

**Figure 8.**
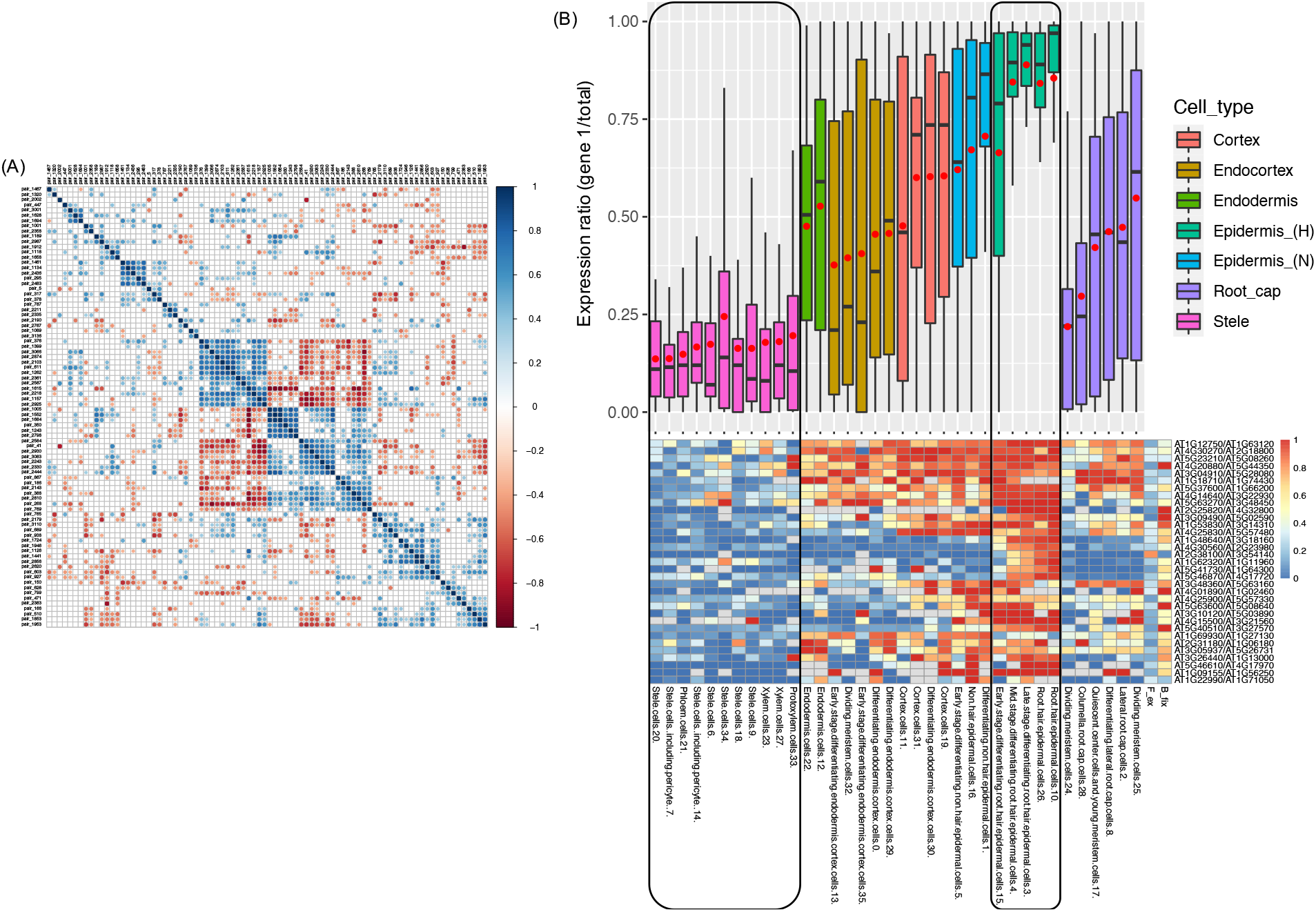
Evidence for concerted divergence among alpha homoeologues. **(A)** Correlogram of pairwise correlation coefficients of expression ratios for all 84 alpha Class 4 gene pairs. Each row or column represents a single gene pair. Significant positive pairwise correlations (scA from both pairs show similar patterns of dominance/non-dominance) are shown in blue, and significant negative correlations (scA from one pair shows a similar pattern as scB from the other pair) are shown in red. Nonsignificant correlation coefficients are shown in grey. A cluster of 33 significantly correlated pairs is visible in the middle of the plot, in which each pair has diverged in expression in a correlated manner with each other pair, suggesting concerted divergence. **(B)** Boxplot and heatmap showing the distribution of expression ratios (scA/total) for the 33 gene pairs comprising the largest cluster of alpha gene pairs exhibiting putative concerted divergence. Expression ratios are shown for each of the 36 RCCs (cell states), and these are clustered and color coded by membership in the 9 superclusters (cell types). Red points in the boxplot indicate mean values.

Within the 33 pair cluster, paralogues have diverged in expression such that one copy of each pair is preferentially expressed in the stele and the other copy is preferentially expressed in the epidermis (Figure 8B). A continuous gradient of bias was evident in intervening layers. The stele-dominant copy of each pair is also dominant in columella cells and one population of meristematic cells in the root cap, whereas the epidermis-dominant copy is weakly dominant in a second population of dividing meristematic cells.

The 66 genes in the cluster are not enriched for any GO terms or protein domains, though several genes are involved in calcium signaling, and there are two pairs of MYB transcription factors. Similarly, neither set of 33 genes in the two separate co-expressed networks is enriched for GO terms, and both homoeologues have equivalent annotations in most cases. Thus, there is no obvious functional differentiation discernable at the level of gene ontology between homoeologues in the two networks.

Of the 13 genes in the epidermal network that have protein-protein interactions annotated in the InTact database (https://www.ebi.ac.uk/intact/), there are two genes whose proteins interact directly: a MYB transcription factor encoded by AT2G31180 and a calcium-sensing calmodulin protein encoded by AT4G14640. A third gene (AT4G30560) encodes a calmodulin-regulated ion channel that interacts indirectly with the calmodulin protein via a protein kinase intermediary (AT4G04570), suggesting a gene module involved in calcium-dependent ion trafficking.

In contrast, none of the corresponding homoeologues in the stele network are annotated as interacting with each other, and no genes from the epidermally-biased co-expression network directly interact with genes from the stele-biased network. Collectively, these observations suggest that the two homoeologous networks have diverged in concert, both spatially and functionally.

### Correspondence of RCC expression classes with functional divergence of paralogues

Expression and function are generally synonymized in discussions of gene evolution (e.g., Panchy et al., 2019). For gene pairs with biased paralogue expression profiles, we therefore assessed the correspondence of different expression patterns with other measures of functional differentiation.

Hanada et al. (2009) categorized 492 Arabidopsis gene pairs as showing low, medium, or high morphological diversification on the basis of knockout phenotypes for one or both paralogues. Of these, only 100 were included among the Wang et al. (2013) duplicate pairs, of which one or both copies were expressed in roots for 96. Despite the small number of pairs in both data sets, the distribution of pairs exhibiting or not exhibiting some degree of morphological variation differed significantly by expression class (χ^2^ = 15.2, p < 0.01). Unlike all other Classes, the majority of Class 2 pairs (no biased paralogue expression) exhibited no morphological divergence, and no Class 2 pairs exhibited high morphological divergence (Figure 9A). If this small sample is representative, the results suggest that pairs of genes with unbiased paralogue expression can tolerate the loss of expression from one paralogue, further suggesting that the two paralogues are not currently maintained either by dosage constraints or by essential functional differences between the two paralogous proteins.

**Figure 9.**
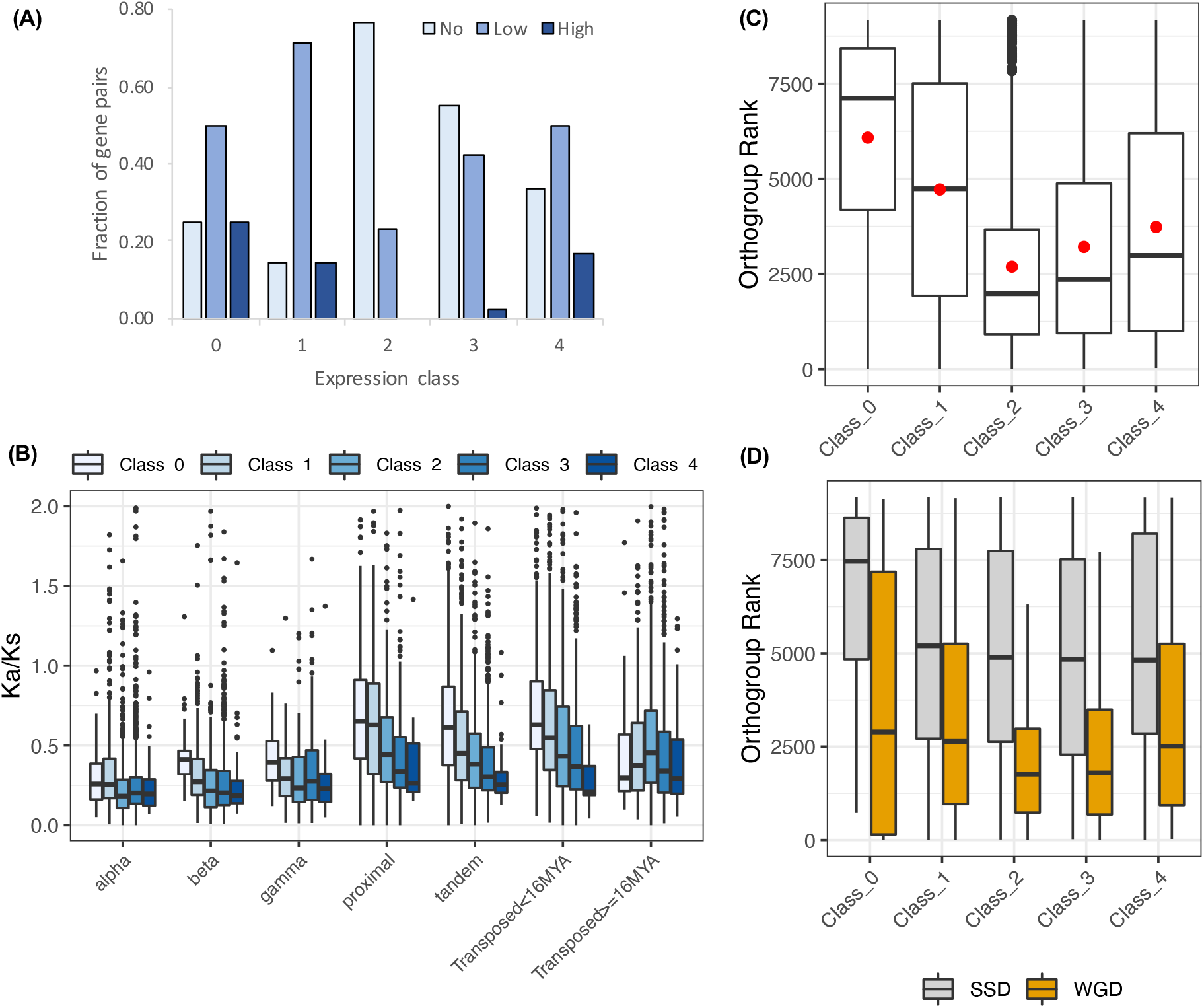
Evidence for functional divergence and gene dosage sensitivity. **(A)** Levels of morphological divergence (Hanada et al., 2009) by expression class. **(B)** K_a_/K_s_ by duplication mechanism and expression class. K_a_/K_s_ values were cut off at 2 for clarity of presentation, but extended beyond this value in all cases. **(C)** Correspondence of expression classes with dosage sensitive gene families (“orthogroups”: Tasdighian et al., 2017). Lower orthogroup rank indicates greater overall dosage sensitivity. Orthogroup rank by expression Class for 5,387 duplicate pairs. **(D)** Orthogroup rank by expression Class separately for WGD and SSD duplication types.

In contrast to Class 2, only around 10% of Class 1 pairs were in the no morphological divergence category. This is likely due to functional differentiation between a root-expressed paralogue and a paralogue expressed in a different context. Classes 3 and 4 were intermediate between Classes 1 and 2 (Figure 9A). Like Class 1, Classes 3 and 4 both partition paralogue expression, but in these Classes partitioning is within the root, between different RCCs. Shared root expression could suggest less functional differentiation between paralogues; moreover, many pairs have both paralogues expressed in RCCs other than those fixed for one paralogue, making those RCCs more like Class 2 pairs. The higher proportion of Class 4 pairs showing some degree of morphological differentiation suggests that balanced fixation--the near-exclusive use of a different paralogue in different RCCs--may be indicative of a shift in function between paralogues.

The ratio of replacement to silent nucleotide substitutions (K_a_/K_s_) is understood to be a measure of functional divergence. For all duplication types and expression Classes, K_a_/K_s_ distributions were strongly skewed, with median and mode less than 1.0 (Figure 9B; Supplemental Table 6), indicating that most gene pairs are evolving under purifying selection, as expeted. As reported by Qiao et al. (2019) for 141 phylogenetically diverse plant genomes, average K_a_/K_s_ varied with duplication type, with WGD classes showing lower K_a_/K_s_ than tandem, proximal, and young transposed pairs. Variances among both duplication types and expression Classes differed significantly (p < 0.001), suggesting differences in the number of pairs showing positive selection or containing an expressed pseudogene. Of the classes, Class 1 had the highest standard deviation in total and for all duplication types except proximal SSDs, suggesting that this Class includes either pseudogenes, genes under positive selection, or both. Class 4 had the smallest standard deviation (Supplemental Table 6). The large variance for Class 1 is consistent with its having the highest percentage of morphologically divergent paralogue pairs as classified by Hanada et al. (2009).

The fact that Class 4 has the lowest average K_a_/K_s_ and the smallest variance around the mean seems to indicate uniformly strong purifying selection. This is counterintuitive with the notion that Class 4 paralogues have diverged functionally, suggesting that they are diverging in their promoters (sequence and/or accessibility) and not in their amino acid sequence.

### Dosage sensitive gene families

Tasdighian et al. (2017) ranked 9,178 core gene families (“orthogroups”) across 37 angiosperm based on how strongly paralogues have been retained in duplicate following WGD vs. SSD; gene families with high levels of WGD retention and minimal SSD retention are defined as “reciprocally retained”, and strong reciprocal retention is considered a hallmark of selection to maintain dosage balance. 6,308 Wang et al. (2013) pairs occur among these 9,178 orthogroups; 853 of these pairs had paralogues assigned to two different orthogroups, and an additional 68 were not assigned to expression Class because one gene model was obsolete; these were excluded, leaving 5,387 pairs with one or both members assigned to an orthogroup and Class. We divided the 5,387 pairs by expression Class and determined the “orthogroup rank” as in Tasdighian et al. (2017; low rank = high reciprocal retention; Figure 9C). Class 2 pairs had the lowest rank, significantly lower than other Classes (Class 2 vs. Class 3: z = −2.22, p = 0.029; Dunn’s test of multiple comparisons with Benjamini-Hochberg correction; all other comparisons significant at p ≤ 0.0001), which is consistent with the expectation that if dosage balance is responsible for preserving both paralogues, neither paralogue is likely to be strongly biased in its expression. Other Classes showed less evidence of dosage-based constraints, with Class 1 pairs showing the least effect (highest orthogroup ranks), consistent with its paralogues being expressed in different organs. Although Class 3 and 4 pairs have some RCCs fixed for one paralogue, other clusters have both paralogues expressed, so it is possible that dosage balance is necessary in some cell types but not in others, where divergence of expression pattern (“function”) is occurring.

Dividing the 5,387 pairs into WGD and SSD duplication types (Figure 9D) showed the expected result that SSD pairs are less affected by dosage constraints than are WGD pairs. Alpha WGD duplicates showed the same overall pattern of Classes as did the full dataset and the WGD duplicates combined (data not shown).

### Expression patterns of duplicate pairs by subgenome

Schnable et al. (2012) assigned *Arabidopsis* homoeologues from 817 alpha WGD pairs to putative *Arabidopsis* subgenomes, and classified the two homoeologous subgenomes as A vs. B, with A being the dominant subgenome, characterized by lower rates of gene loss and higher expression of remaining genes. We compared subgenome assignments with our scA/scB classification based on expression of Wang et al. (2013) alpha WGD homoeologues in RCCs, and found representation of pairs from all our classes, with 9 Class 0 pairs, 82 Class 1 pairs, 240 Class 2 pairs, 377 Class 3 pairs and 23 Class 4 pairs (86 pairs from Schnable et al. (2012) were not included in the Wang et al. (2013) assignments). Class 0 and Class 2 pairs by definition have no bias detected in our data, while Class 1 may have bias, and Classes 3 and 4 must have at least one biased context. Of the 453 pairs in the Schnable et al. (2012) assignments for which we did see bias in one or more contexts, the scA/scB assignments based on dominance observed in our expression data were consistent with the A/B subgenome dominance assignments 261 times and inconsistent with them 192 times. Of the three Classes, Class 1 was most concordant, with approximately 4 times the number of pairs agreeing than disagreeing, while Classes 3 and 4 both had roughly only 1.25 times more cases of agreement than disagreement. The average F_ex_ value for pairs in agreement with the subgenome assignments was 0.36 compared to 0.31 for those in disagreement.

We also asked if the 33 alpha pairs representing a putative example of concerted divergence (Figure 8) have partitioned expression by subgenome. The stele-dominant set includes eight homoeologues from subgenome A and five from subgenome B, and the epidermis-dominant set includes five homoeologues from subgenome A and seven from subgenome B. 41 of the 66 genes in the two sets genes were not assigned to subgenome by Schnable et al. (2012). This lack of subgenome assignment for most genes makes it difficult to assess patterns of subgenome partitioning, but the mixed representation in each pathway suggests that the two sets most likely diverged in concert after the alpha WGD, enlisting genes from both subgenomes, rather than in the progenitors of the polyploid.

### Expression differences within cell types

The 9 superclusters identified by Ryu et al. (2019) in some cases aggregated known cell types (e.g., Ryu supercluster 1, in which cells of the quiescent center are grouped with root cap cells; also Ryu superclusters 0, 2, and 5). These presumably artificial superclusters were disaggregated in the 36 RCC data studied here (e.g., Figs. 2, 3, 6). This resulted in the identification of some alternate fixation patterns between RCCs of the Ryu et al. (2019) superclusters. For example, there were 32 alternate fixations within Ryu supercluster 5, with protoxylem cells being particularly differentiated in their expression pattern relative to other cell types grouped in the same supercluster.

Ryu et al. (2019) studied the developmental trajectories of root hair cells, as well as other cell types, and those cell developmental states are also included in the 36 RCC data, allowing us to search for differential responses to duplication among developmental states of more confidently identified cell types: non-hair epidermal cells (three RCCs of supercluster 3), root hair epidermal hairs (three RCCs of supercluster 4), late stage and mature cortex cells (four RCCs of supercluster 6), mature endodermis cells (two RCCs of supercluster 7), and a second root hair supercluster (two RCCs of supercluster 8). No cases of alternate fixation were found among states belonging to superclusters 4, 7, or 8, but a beta pair (AT3G13750/AT5G56870) was found to be alternately fixed for states within both superclusters 3 and 6; the alpha pair AT3G18950/AT1G49450 (row 12 in Figure 2) and the young transposed pair AT5G39580/AT5G64100 were found to be alternately fixed for states within supercluster 6.

In addition to the small number of reciprocal shifts in paralogue bias (alternate fixation) among states of the same cell type, there were many other instances where the expression patterns of paralogue pairs were not homogeneous across states of a given homogeneous cell type (e.g., rows 1, 2, 7, 8, 11, and 12 in Figure 2). For example, there were 87 alpha WGD pairs that had no expression of either paralogue in one RCC of non-hair epidermal cell supercluster 3, fixation of one paralogue in a second RCC of the supercluster, and balanced expression of both paralogues in the third RCC (Supplemental Table 7). Overall, for supercluster 3, 35% of alpha WGD pairs (1,119/3,181) had non-homogeneous expression across its three RCCs; for all duplication types, the average was 30% for this supercluster. In all five of these superclusters, the three WGD classes and the older transposed duplication class had higher percentages of non-homogeneous expression; the actual percentages were roughly correlated with the number of RCCs in a supercluster--finer division resulted in greater heterogeneity--but superclusters with the same number of RCCs differed from one another (e.g., superclusters 3 and 4, each with three RCCs, had overall heterogeneity percentages of 30% and 23.3%, respectively), suggesting cell type-specific patterns of paralogue expression during differentiation.

### Shared and unique transcriptomes of cell clusters

The 22,669 genes that are expressed in at least one of the 36 RCCs include 7,653 genes, comprising 33.8% of the overall root transcriptome and well over half of the transcriptomes of some RCCs, that are expressed in all 36 root cell clusters (RCC-u genes; Figure 4). Among genes expected to belong to the RCC-u class are genes that are required in all root cells for root-specific functions, and genes that are expressed in all cells of the plant (both root and non-root). Cheng et al. (2017) identified a set of 4,577 genes expressed in all 11 Arabidopsis tissues. 3,505 RCC-u genes were among these 4,577 ubiquitously-expressed genes. Thus, most RCC-u genes are not root-specific, a fact further emphasized by the expression of over 99% of RCC-u genes in at least one non-root SRA dataset (data not shown).

RCC-u genes are an interesting class to consider because their presence in all RCC transcriptomes suggests that their expression is indispensable in all root cell types and states, and perhaps in all cells, and further suggests that, initially after any duplication—whether WGD or SSD—both paralogues were expressed in all RCCs (both paralogues were RCC-u). We therefore explored the degree to which both paralogues of RCC-u gene pairs also retained their status as RCC-u genes.

Among the 7,653 RCC-u genes are 4,538 genes belonging to 4,639 Wang et al. (2013) gene pairs. 40 of these gene pairs include one gene that is RCC-u and one that is an obsolete gene model; consequently, these pairs are not assigned to an expression class, leaving 4,599 Wang et al. (2013) RCC-u gene pairs assigned to an expression class. Only 1,552 pairs retained both genes as RCC-u, and among these, only 847 pairs did so without significant expression bias (Class 2; Table 3). Thus, both copies retain the putative ancestral expression profile for only 18.4% (847/4,599) of RCC-u pairs. 705 pairs retained both genes as RCC-u but showed biased expression in one or more RCCs (Classes 3 and 4). The remaining 3,047 pairs include one copy that is RCC-u and one that is not, indicating changes in expression in at least one copy from the ancestral profile. For 327 pairs, these shifts were subtle, as we did not detect significant bias (Class 2). In the remaining cases (2,720 pairs), the shift from the hypothesized ancestral condition of both paralogues being RCC-u was more dramatic, including the 505 pairs for which the second paralogue was not expressed in any root cell type (Class 1; Table 3).

**Table 3.** Breakdown by expression Class of 4,599 Wang et al., (2013) paralogue pairs ubiquitously expressed in root cell clusters (RCC-u gene pairs).

| Class | Count | RCC-u mixed | RCC-u both |
| --- | --- | --- | --- |
| 0 | 0 | 0 | 0 |
| 1 | 505 | 505 | 0 |
| 2 | 1,174 | 327 | 847 |
| 3 | 2,780 | 2,126 | 654 |
| 4 | 140 | 89 | 51 |

These counts are further broken down by duplication mechanism in Supplemental Table 8. For alpha duplicates, there were 1,685 pairs with one or both genes RCC-u. Of these, in 294 pairs both homoeologues were RCC-u but show significant bias in at least one cluster (17.4%; Classes 3 and 4), and 949 pairs (56.3%) have only one RCC-u copy with varying degrees of bias in clusters where both homoeologues are expressed (Supplemental Table 8). Conversely, 442 pairs (26.2%) show no evidence for shifts from the ancestral state (both copies are RCC-u with no bias [Class 2]). This is the highest fraction of pairs with both copies retaining the ancestral expression pattern of any duplication type. Gamma and beta WGD duplicates exhibited the next highest fractions (16% and 15.1%, respectively), and the SSD duplicates had the lowest. This suggests that WGD duplicates are more constrained to retain their ancestral expression patterns, perhaps due to dosage constraints.

Representation in the RCC-u class itself also varied with duplication type, with over-representation of WGD duplicates expressed across all RCCs relative to the total fraction of WGD duplicates in the genome (Supplemental Figure 4). In contrast, SSD duplicates, other than older transposed duplicates, were under-represented relative to their representation in the genome as a whole, more like genes lacking a duplicate in the Wang et al. (2013) set and for which one paralogue has presumably been lost after duplication (“singletons” in Supplemental Figure 4).

Evolutionary shifts in expression of paralogue pairs were also observed among the remaining 59.6% of pairs comprising the cumulative root transcriptome that are expressed in 1-35 RCCs (non-RCC-u pairs). Their lack of ubiquitous expression in root cell types/states makes it more difficult to determine whether expression of a given gene in a particular RCC represents the ancestral state or the derived state, and thus to hypothesize the ancestral condition for a paralogue pair with only one member expressed in an RCC. If we assume that at the time of a duplication, both paralogues retained the expression pattern of the single copy gene progenitor (see below for discussion of this assumption), then pairs with unbiased expression of both paralogues or neither paralogue in a given RCC are most readily explained as retaining the expression state of their single copy progenitor, since two independent gains or losses of expression would need to be hypothesized otherwise.

Pairs with only one paralogue expressed in any RCC (Class 1), or with biased expression of one paralogue (Classes 3 and 4), could equally parsimoniously be inferred to have gained or lost expression of one paralogue in RCCs for which the pair shows fixation. In either case, however, an evolutionary shift in expression from the inferred ancestral condition is involved. The total number of RCCs showing fixation of a paralogue pair in the non-RCC-u class (i.e., the number of red or blue cells in a heatmap of non-RCC-u pairs, similar to the example shown in Figure 2) was 22,897; this was 45% of the total number of number of heatmap cells (non-RCC-u paralogue pairs × 36 RCCs; Supplemental Table 9).

Because we were interested in exploring differential responses of different cell types/states to different types of gene duplication, we calculated a “paralogue expression retention” (PER) index, defined as the number of pairs expressing both paralogues in a given RCC divided by the number of pairs with at least one paralogue expressed in the RCC, and compared this value across RCCs for all pairs except those for which both paralogues were classified as RCC-u. There was considerable variation in PER by both RCC and duplication type, with WGD and older transposed duplicate pairs showing higher rates of paralogue retention (Figure 10A-B). Variation in PER across RCCs was likely due, in part, to differences in sequencing depth per RCC (RCCs with fewer total reads are more susceptible to dropout). On the assumption that sequencing depth has the same relative effect on PER for different duplication mechanisms in all RCCs, we adjusted for differences in sequencing depth by calculating the differential retention ratio, PER_wgd_/PER_ssd_ for each RCC. Because of our specific interest in the alpha WGD we also calculated PER_alpha_/PER_tandem_ (Figure 10C). These two ratios were similar (Figure 10D). Variation across RCCs is likely due to differences in cell biology that reflect the degree to which the genes whose expression comprises the transcriptome are dosage sensitive; higher PER of WGD pairs is expected given their greater representation among dosage sensitive gene families (Tasdighian et al., 2017; Qiao et al., 2019).

**Figure 10.**
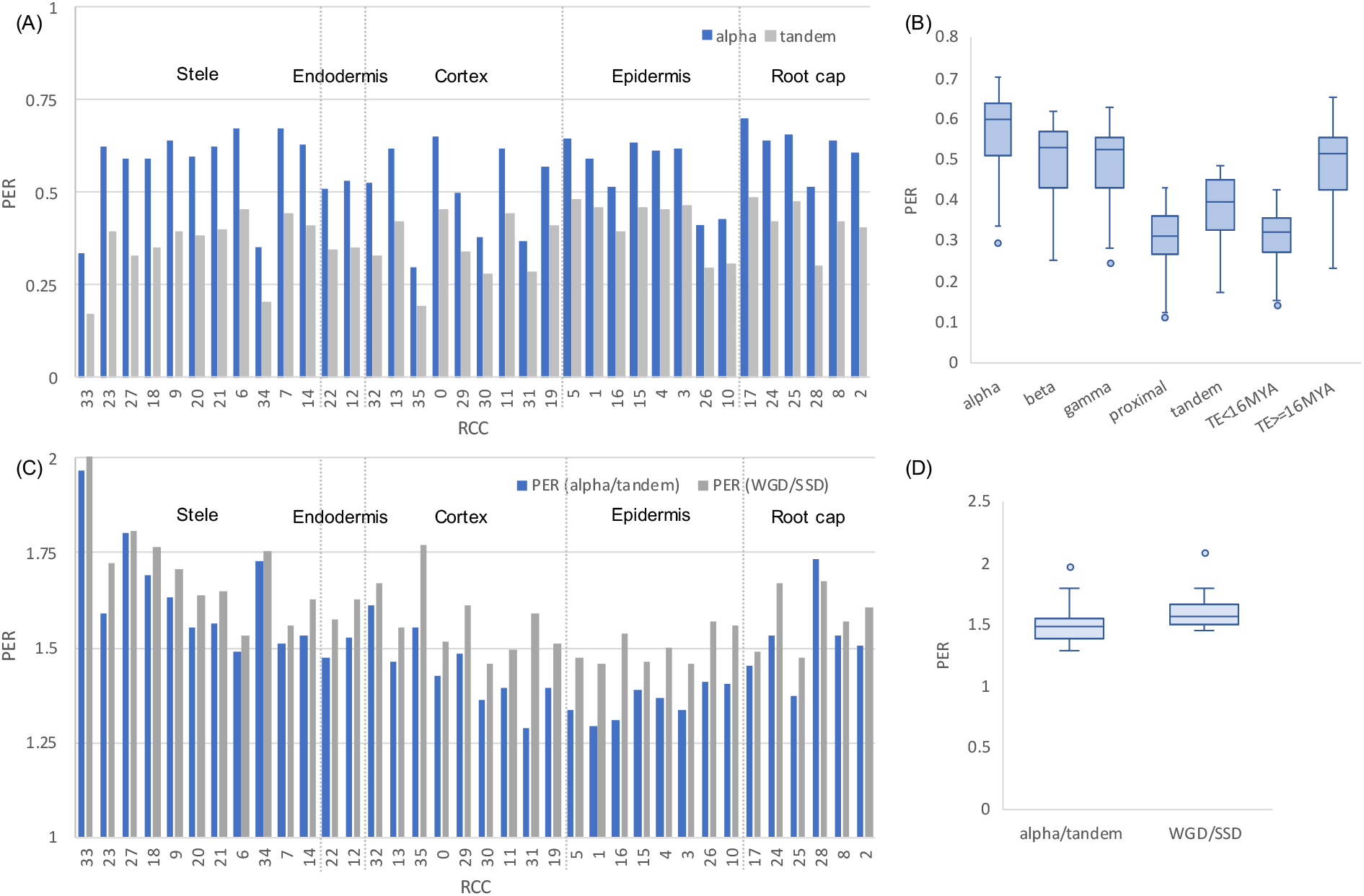
Paralogue retention ratio (PER) by RCC and duplication mechanism. **(A)** PER for alpha and tandem duplicates by RCC. The grouping of RCCs by cell type is indicated with dashed lines. **(B)** PER distributions across RCCs by duplication type. **(C)** Differential retention ratios for WGD/SSD and alpha/tandem duplication types. **(D)** Distribution of differential retention ratios (alpha/tandem and WGD/SSD).

## DISCUSSION

The primary goal of this study was to explore how knowledge of gene expression at the single cell level can elucidate the process of evolution following polyploidy, with a secondary emphasis on single gene duplications. The basic premise is simply an extension of the justification for all single cell biology: that the cell is a more basic level of organization than tissues or organs, and that aggregating different cell types and cell states in any analysis obscures patterns that may be of interest (e.g., Efroni and Birnbaum, 2016; Libault et al., 2017). This is by no means a new idea, but it has not been applied to polyploidy, with the notable exception of the elegant and extensive work on cotton fibers, which are single cells of great economic importance and an excellent system for studying development at the cellular level (Shi et al., 2006; Gou et al., 2007; Taliercio and Boykin, 2007; Hovav et al., 2008a, b; Al-Ghazi et al., 2009; Rapp et al., 2010; Wang et al., 2010; Nigam et al., 2014; Yoo and Wendel, 2014; Hu et al., 2015; Tuttle et al., 2015; Bao et al., 2019; Gallagher et al., 2020). The recent spate of papers on *Arabidopsis* root biology at single cell resolution (Jean-Baptiste et al., 2019; Ryu et al., 2019; Shulse et al., 2019; Denyer et al., 2019; Zhang et al., 2019) provides comparable information from individual cells (rather than pooled cells) from multiple cell types and their developmental states. Coupled with the longstanding interest in the polyploid nature of the *Arabidopsis* genome that has led to the classification of its duplicated genes by mechanism of duplication (e.g., Wang et al., 2013; Hao et al., 2018; Qiao et al., 2019), this has provided us with the opportunity to explore the evolution of gene expression in over 11,000 paralogous pairs at a finer scale than has previously been reported. We observed numerous cases of greatly biased paralogue expression among cell types and states of the root, in several cases involving alternate bias of the two paralogues (Figure 5–7). We identified over 7,000 genes ubiquitously expressed in root cells, which afforded us strong hypotheses of ancestral expression, and allowed us to infer extensive expression shifts since paralogue or homoeologue divergence.

There are two perspectives to consider concerning the response to genome duplication: that of the cell and that of the gene. From the standpoint of the cell, the issue is one of function—how does a cell type, optimized functionally in the course of hundreds of millions of years of evolution, adjust to the opportunities and challenges of having its gene repertoire suddenly duplicated? Arabidopsis allows us to address this question after millions of years of evolution during which the genome has fractionated, retaining only a subset of its genes (e.g., Cheng et al., 2018; Emery et al., 2018; Liang and Schnable, 2018). How are these retained duplicates deployed in a given cell type? In contrast to the work on cotton fibers, the *Arabidopsis* root single cell transcriptome data provide us with 36 cell clusters comprising several cell types with up to several states each. Do all root cell types respond similarly in terms of degree to which they deploy genes that remain duplicated?

From the complementary perspective of the gene, questions involve the mechanisms that have led to the preservation of only a fraction of the genes from two complete homoeologous genomes. This fraction of the genome includes, among other things, genes maintained in duplicate because of dosage constraints, that need not involve any new function or expression pattern (e.g., Freeling, 2009; Birchler and Veitia, 2014; Tasdighian et al., 2017; Defoort et al., 2019). Other pairs of homoeologues are retained, through various different mechanisms, because their functions have evolved to make both copies indispensable (Panchy et al., 2016; Qiao et al., 2019). The fine-grained perspective of single cells and cell types allows us to ask whether there is evidence for differentiation of function or expression among cell types of a single organ, the root, much as Adams et al. (2003) addressed this issue at the level of the component organs of the flower. At an even finer level, are different homoeologues expressed in different cells of the same cell type or cell state? And if expressed within the same cell, do homoeologues participate in different networks, perhaps as a result of concerted divergence (Blanc and Wolfe, 2004)?

### The limitations of our single cell data

Because all phenotypes originate at the cellular level (Lynch and Trickovic, 2020), it seems reasonable to assume that the biologically relevant unit of analysis for studying the evolution of duplicate gene expression is the individual cell. Each gene is transcribed in the nucleus of a single cell, though its mRNA or protein product may be mobile and function elsewhere, and so it is in individual cells that novel gene expression patterns occur that are central to duplicate preservation by mechanisms such as subfunctionalization and neofunctionalization. It is also in individual cells that multimeric complexes are assembled and gene regulatory networks operate, so it is at the cellular level that the sensitivity to dosage hypothesized to preserve genes in duplicate following WGD events manifests itself (Papp et al., 2003, Freeling, 2009; Birchler and Veitia, 2010, 2012, 2014; Song et al., 2020).

But understanding the expression of a pair of paralogues requires first understanding the expression of individual genes, and this is not a simple matter. In contrast to early concepts of gene regulation, expression is stochastic and noisy, with bursts of transcription interspersed with inactivity, and the levels of transcripts, and thus proteins, depend on complex dynamics of expression and degradation (Araujo et al., 2017; Nicholson, 2019; Tunnacliffe and Chubb, 2020). This complexity has not been incorporated into models that seek to explain duplicate gene retention and loss. As an example, for a protein encoded by a pair of paralogues and for which dosage balance maintenance is important (e.g., as part of a multi-subunit complex), it is the sum of the expression of both paralogues that is critical; the numbers of transcripts produced from either paralogue is less important in the short term, but potentially dictates whether one paralogue will ultimately be lost to the process of fractionation (Gout and Lynch, 2015). This revised version of the duplication-degeneration-complementation (DDC) model (Force et al., 1999) involves gradual evolution of stochastic differences in paralogue expression, but it is now clear that stochastic differences are part of the transcriptional process itself, and that these also can play a role in promoting the preservation of duplicate genes (Rodrigo and Fares, 2018; Chapal et al., 2019).

A particular scA/scB transcript ratio for a paralogue pair in a cell cluster could be due to all cells expressing that ratio (rheostat-like control of expression); to an on/off mechanism, with subsets of the cell population expressing scA and scB, creating the scA/scB ratio by modulating the relative number of cells expressing each gene (Nicholson, 2019); or to more complex modes of transcriptional control (Tunnacliffe and Chubb, 2020). For polyploids, particularly allopolyploids that combine diverged genomes, there exists the possibility that a single cell cluster might include individual cells that are differentiated in their expression by subgenome. This seems unlikely, given the integration observed for higher level phenotypes--an allopolyploid individual is not a mosaic for the two different floral morphologies of its progenitors, for example, but instead integrates the developmental gene networks of its progenitors to produce a distinctive flower. But given the idea that those gene regulatory networks can diverge from one another (Blanc and Wolfe, 2004), which we see here among root cell clusters (Figure 7), it is worth looking for evidence of higher-order independence of genome expression, and the level of the cell is a natural place to look.

However, measuring transcript amounts in individual cells is not trivial technically, with a tradeoff between number of cells and number of transcripts that can be assayed; in the system used to generate these data, the “dropout” effect is a particular problem, and can distort estimates of transcript numbers, particularly between highly and weakly expressed genes (Bhargava et al., 2014; Lähnemann et al., 2020). Despite being able to analyze only a small number of gene pair x cell combinations with statistical rigor, it appears that most gene pairs express both paralogues in individual cells. However, there do appear to be cases where expression of paralogues is partitioned into discrete cells, and this is more often true of gamma WGD homoeologue pairs than for pairs created by other types of duplications (Table 2). As single cell technology improves, it will be interesting to look at larger numbers of cells for any evidence of differentiation by subgenome. As the challenges of dealing with dropout are overcome (Lähnemann et al., 2020), It should also become possible to quantify the extent of deterministic (i.e., rheostat) vs. stochastic (i.e., on/off) gene regulation by determining if differences in expression among cell types are achieved via by changes in the per-cell expression level or by changes in the fraction of cells expressing the gene.

### Cell types, cell states, and cell clusters

The next highest unit of organization above the cell is the “cell type” and it is at this level that emergent properties of stochastic gene expression within individual cells should manifest themselves. The cells of the root are all derived from the root apical meristem, and thus all are members of the same cell lineage. The basic cell types of the root—xylem, phloem, epidermis, cortex, root hairs, root cap, etc.—are conserved across most plants that have roots, and presumably evolved early in the history of vascular plants (Raven and Edwards, 2001; Kenrick and Strullu-Derrien, 2014; Huang and Schiefelbein, 2015). That cell types exist is taken as a given in the molecular and developmental biology literature, but defining this term is not simple. Although there has been recent theoretical progress in how cell types originate and evolve (e.g., Arendt et al., 2016; Liang et al., 2018; Yuan et al., 2020), Vikaryous and Hall (2006) liken the problem of defining “cell type” to defining “species”, an endless source of controversy in evolutionary biology. All of the recent Arabidopsis root single cell papers (Jean-Baptiste et al., 2019; Ryu et al., 2019; Shulse et al., 2019; Denyer et al., 2019; Zhang et al., 2019) use the term “cell type” but it is not defined in any of them.

Operationally root single cell studies use algorithms to aggregate single cells into clusters of transcriptomically similar cells. These clusters are assigned to known cell types using genes that have been reported to be characteristically—sometimes uniquely—expressed in those cell types, but because such marker genes are not available for all possible cell types and subtypes, the correspondence between root cell clusters (RCCs) and cell types is not perfect. For example, Cluster 1 of Ryu et al. (2019) contains some cells identified as belonging to root cap whereas others come from the quiescent center. In any case, one of the major goals of single cell transcriptomic studies is to identify previously unknown cell types. Such discovery is complicated by the fact that, according to Morris (2019), “cell identity is stable; however, in response to diverse stimuli, the same cell type can exhibit a range of different phenotypes (states).” One of those “diverse stimuli” is development, and the root tips studied in Ryu et al. (2019) allowed them to follow the differentiation from meristematic cells to diverse mature cell types. For our study, 36 root cell clusters (RCCs) were the primary units of analysis, each assumed to be a single cell type and, for a subset of the 9 superclusters of Ryu et al. (2019) that appear to correspond to single cell types, developmental states of a single cell type. These hypothesized relationships are consistent with the nested relationship of the 36 and 9 cluster transcriptomes (Figure 3).

### Expression evolution of alpha WGD homoeologue pairs in root cell types and cell states

For simplicity we assume here, as others have done (e.g., Blanc and Wolfe, 2004; Panchy et al., 2019; but see below) that most alpha WGD pairs initially conserved their ancestral function and partitioned their expression equally between what are now their two homoeologues. However, determining whether the expression of only one paralogue in an RCC is the ancestral or the derived state is not straightforward for most pairs, as no close outgroup is available (e.g., the closest relative to an alpha duplicate pair generally is a beta WGD homoeologue, diverged around 100 MY from either alpha homoeologue; Panchy et al., 2019). Therefore, we focused on gene pairs where at least one member of an alpha homoeologue pair was ubiquitously expressed in all root cell clusters (i.e., pairs that contained at least one RCC-u gene, detectable in all 36 RCCs). We reasoned that these genes were likely to be necessary for the functioning of all cell types and cell states in roots, so that the ancestral condition of the pre-alpha WGD single copy gene was also ubiquitously expressed in root cells.

With these two assumptions, we looked for departures from the ancestral condition, representing shifts in expression following homoeologue divergence from their common ancestor. Out of 1,685 alpha pairs with at least one RCC-u homoeologue, we found 949 with only one RCC-u homoeologue, including 94 pairs with only one homoeologue expressed in roots (Class 1 alpha WGD pairs; Supplemental Table 8). Additionally, we observed 294 alpha pairs with both homoeologues RCC-u, but with biased expression of one or both homoeologues in one or more RCCs (280 Class 3 and 14 Class 4 pairs, respectively). Thus, in 73.7% of RCC-u alpha WGD pairs, at least one member has changed its expression as homoeologues have diverged.

Determining ancestral states by this logic does not apply to non-RCC-u pairs--those with neither member expressed ubiquitously across the 36 RCCs. However, even without being able to hypothesize the polarity of the change, under the assumption that both paralogues initially were expressed in an unbiased fashion in the same cell immediately after their duplication we can assess the amount of change, and found that 72% of alpha WGD non-RCC-u pairs had one homoeologue uniquely or preferentially expressed in at least one RCC. High levels of expression differentiation between homoeologues are not unexpected. For example, in the older (ca. 60 MY) polyploid, *Gossypium raimondii*, over 90% of homoeologues show evidence of sub- or neofunctionalization (Renny-Byfield et al., 2014).

Determination of expression patterns at the level of root cell types/states also provides information on mechanisms of gene retention. The 590 alpha WGD RCC-u gene pairs that express both copies without strong bias (Class 2 RCC-u pairs; Supplemental Table 8) are candidates for preservation by dosage balance constraints (Papp et al., 2003; Coate et al., 2016; Tasdighian et al., 2017; Song et al., 2020). More broadly, all Class 2 pairs, including those that are not expressed in all 36 RCCs (non-RCC-u) are also candidates for preservation by dosage balance. Consistent with this, Class 2 alpha WGD genes showed less evidence of functional divergence (Figure 9A) than did other Classes, were more likely to belong to dosage-sensitive gene families (Figure 9C-D), and, as expected, showed less evidence of positive selection than did SSD pairs or alpha pairs in other expression classes (Figure 9B).

Alpha WGD pairs belonging to Classes 3 and 4 provide examples where homoeologues could be maintained by sub- or neofunctionalization. The polarity of expression shifts of the 1,570 alpha WGD Class 3 pairs is unknown, but each includes at least one RCC where there is novel bias subsequent to the divergence of the two homoeologues. This could occur by enhanced expression or recruitment of one homoeologue, the latter being consistent with neofunctionalization. Alternatively, bias could be achieved by diminished expression of the weakly-expressed homoeologue (subfunctionalization). For Class 3 pairs we do not know the cell-level expression site of the scB paralogue, so it is unknown whether scB is predominantly expressed in any non-root cell type. For Class 4 pairs, however, there is alternate fixation of paralogues among the 36 RCCs, providing self-contained examples consistent with subfunctionalization of 88 alpha WGD pairs just within root cell types, novel information not available from bulk tissue samples (Figure 7).

### Response of cell types to the alpha WGD event

Immediately following each of the three polyploidy events detectable in genomes of most Brassicaceae, the root apical meristem of the neopolyploid plant successfully utilized its suddenly multiplied genome to carry out its programmed functions, generating the various cell types in their characteristic developmental progression to form a functional root. Each cell type produced from the meristem must also have functioned well enough, through all of the developmental states leading to its mature condition, to allow the plant to survive.

Beginning in the earliest generations--perhaps even immediately, as a response to genomic shock (McClintock, 1984)--and continuing through the ongoing process of diploidization, the combination of neutral and selective forces that operate on variation beginning at the cellular level (Lynch and Trickovic, 2020) led to changes in expression patterns of gene pairs that remained in duplicate, in turn setting the stage for retention of some homoeologues and loss of others (fractionation). These events, along with speciation and individual gene duplication, produced the *Arabidopsis thaliana* genome, with its current subgenome structure (Schnable et al., 2012).

Individual *Arabidopsis* root cell types and states comprising the 36 RCCs evolved their current expression patterns over 30-40 MY of speciation, divergence, and diploidization since the alpha WGD event, each expressing a subset of the homoeologues retained in the genome. How have these different cell types/states responded to the opportunities and challenges of a doubled genome, specifically by expressing only one copy or expressing both homoeologues? 73.7% of the 1,685 alpha WGD RCC-u gene pairs showed evidence of evolutionary shifts in expression in at least one cell type or state (Supplemental Table 8), and many pairs showed such shifts in many or all RCCs (e.g., by definition a Class 1 RCC-u pair shows an expression shift in each of the 36 clusters). Overall, we found that for both RCC-u and non-RCC-u alpha WGD pairs, over one third of all cell type/state x homoeologue pair combinations for which statistically robust measurements could be made showed evidence of expression shifts (Supplemental Table 9).

Our results show that each cell type/state has had its own characteristic response to gene and genome duplication. For example, excluding RCC-u genes, approximately 60% of alpha WGD pairs had both homoeologues expressed per cluster on average across all 36 RCCs, but in individual cell types/states this value varied from around 30-70% (Figure 10A-B). Moreover, the propensity to express both WGD homoeologues as opposed to both SSD paralogues varied among cell types/states. Specifically, across all cell types/states and gene pairs, both homoeologues of alpha WGD RCC-u pairs were expressed about 1.5x more frequently than were both paralogues of tandem SSD pairs, but varied from 1.3x to nearly 2x (Figure 10C-D). Variable responses of different cell types/states is also seen clearly in levels of biased expression of the two paralogues of duplicate pairs in the 36 RCCs, with biased expression of alpha WGD pairs averaging around 40% over all RCCs, but varying from less than 25% to nearly 45% showing bias depending on the cell type/state (Figure 6). We attribute this to functional differences between cell types/states being reflected in the kinds of genes being expressed there, which in turn is connected directly to the mechanisms that lead, through expression differences, to differential retention of pairs produced by different types of duplication (Tasdighian et al., 2017; Qiao et al., 2019).

### Assumptions about polyploidy

Using the Arabidopsis alpha WGD as our test case for studying the evolution of polyploid gene expression at the cell level imposes certain limitations that are common to most if not all older (paleo) polyploids (e.g., Doyle and Egan, 2010). The greatest of these is our lack of knowledge concerning the nature of the polyploidy event, and thus we are ignorant of the degree of differentiation of the two homoeologous genomes we are studying millions years after their divergence from one another. The alpha WGD event is widely accepted to have been allopolyploid, based on biased fractionation of the homoeologous genomes of *Arabidopsis* (Schnable et al., 2012; Woodhouse et al., 2014; Cheng et al., 2018).

Allopolyploidy is a composite phenomenon involving the separate processes of genome merger and genome duplication, as well as their interaction. In many if not most cases, it is a process rather than a singular event, with duplication preceding merger or vice versa depending on the pathway, and often involving intermediate states before producing a tetraploid from two diploid progenitors (Mason and Pires, 2015). It is unknown what path the alpha WGD took in merging the genomes of two diverged genotypes that perhaps belonged to what we would today classify as species. This does not change the empirical data on patterns of homoeologous gene expression that we report here, but it does leave open the question of whether most differences in expression originated at the time of diploid progenitor speciation or after genome merger and doubling (in whatever order these occurred).

Based on the phylogenetic distribution of taxa with and without the alpha WGD, Edger et al. (2018) bracket the date of the allopolyploidy event at around 30-40 MYA. Thus, the progenitor species genomes had as much as 10 MY to differentiate prior to being brought together in the same nucleus—nearly twice the divergence time of chimpanzees and humans (Antonelli et al., 2017). It is unlikely that their roots were identical, any more than one would expect their floral or leaf morphologies to be identical, but although root architecture and anatomy of angiosperms can vary greatly, including in the number and types of cells/tissues (e.g., endodermis in grasses: Enstone et al., 2002; Pacheco-Villalobos and Hardtke, 2012), it seems likely that congeneric species would possess the same root cell types, and that cell types would each pass through similar developmental states in each species. The amount of orthologue divergence in the two progenitor species should be tied intimately to the evolution of the functions of the cells in which they are expressed. Given the presumably conservative nature of cell type evolution, orthologue expression evolution thus might also be expected to be relatively conservative. Even so, the same cell type can differ considerably even among congeneric species both in morphology and transcriptomically, as is true of fiber cells in the diploid species whose genomes merged to form tetraploid cotton (Yoo et al., 2013; Gallagher et al., 2020). Thus, differences in the expression patterns of alpha WGD homoeologues may be due to “parental legacy” (Buggs et al., 2014; Steige and Slotte, 2016) and not solely to genome duplication.

Nevertheless, it seems likely that, on average, more of the differentiation of alpha WGD homoeologues has occurred subsequent to genome merger, for three reasons. First, the time since polyploidy is at least 3x longer than the time since speciation prior to allopolyploidy (Edger et al., 2018), so even if homoeologue divergence has been proportional to time, more differentiation would be expected to have taken place in the last 30 MY than in the first 10 MY since progenitor speciation. But, second, genome evolution after allopolyploid merger and duplication is thought to be far from the clock-like divergence that might be expected after speciation, beginning with an initial “genomic shock” phase of rapid genetic and genomic change, followed by diploidization that returns the allopolyploid to a more conventional evolutionary rate (Wendel, 2015). Finally, orthologues generally evolve conservatively with respect to function and expression pattern, at least relative to paralogues (Gabaldon and Koonin, 2013).

For any pairs in which parental legacy explains differences in homoeologue expression, the effect on our assumptions is that this pushes the ancestral state back to the ancestor of the diploid progenitors of the polyploid, so that the “duplication event” for a pair of homoeologues is speciation, a process that, like polyploidy, is a whole genome phenomenon rather than a gene-by-gene event. In theory, orthologues are formed simultaneously at each locus by speciation, though of course it is an open question when, precisely, species actually form (De Queiroz, 2007). The ancestral condition of a pair of orthologues is like that of a pair of paralogues formed by a tandem duplication: In both cases the products are expected to be expressed at the same level and in the same cellular context.

### Conclusions and future prospects

The ability to assay transcription at the level of cell types and their developmental states permits elucidation of gene expression patterns at a very fine scale, and although the results we present here largely fall short of true single cell transcriptomics due to the sparsity of the data, it is almost certain that this and other current technical challenges will be overcome (Lähnemann et al., 2020). We can look forward soon to being able to compare the expression of gene pairs not only in root cells, but in all cells of the plant, eliminating the need to compare root single cell data with non-root transcriptomes generated from tissues or whole organs, as we did here.

Our characterization of the patterns of duplicate gene expression has identified a wealth of examples of differential expression of gene pairs among root cell types, the first step in achieving a detailed understanding of such issues as how pairs of homoeologues are regulated in allopolyploids at the level of trans-acting factors and cis-regulatory elements (Hu et al., 2020). Single cell tools already exist for addressing these issues by assaying chromatin accessibility (Farmer et al., 2020) and mapping the epigenomic landscape of the cell (Zhou et al., 2019). For understanding functional differences between paralogues, and for even finer scale “sub-localization” of gene action within cells that may be a driver of paralogue retention (Qiu et al., 2019), a host of other single cell -omics tools exist or are under development (Macaulay et al., 2017; Hasle et al., 2020).

We can learn much from single cell expression studies of a single individual of *A. thaliana*, but it is known that gene expression varies considerably among individuals of this species (Cortijo et al., 2019), and that the response even to autopolyploidy in *Arabidopsis* varies among accessions (Yu et al., 2010), so we look forward to the availability in expression atlases (e.g., Papatheodorou et al., 2019) of single cell data for roots and other tissues of additional individuals, and accessions, and also from other species. Interspecific data would allow us to hypothesize ancestral states with far more confidence, particularly data from Cleomaceae, the sister family to Brassicaceae, whose common ancestor pre-dates the alpha WGD event (Edger et al., 2018).

Moreover, there are fascinating questions about the evolution of cell type transcriptomes that can only be addressed with comparative studies. For example, Liang et al. (2018) describe the phenomenon of “correlated evolution” of transcriptomes, in which transcriptomes of different (non-homologous) tissues of a species cluster together, instead of transcriptomes of homologous tissues in different species clustering together as expected. This is thought to be a consequence of the non-independence of transcriptomes in cell types that share transcription factors and their target genes. It is reminiscent of concerted evolution in gene families and concerted divergence of gene regulatory networks, both of which are phenomena of considerable interest and importance in polyploids (Blanc and Wolfe, 2004; Qiao et al., 2019). Comparative single cell data from other *Arabidopsis* species, including from recently formed allopolyploids in this excellent model system, hold much promise for addressing these and other questions.

## AUTHOR CONTRIBUTIONS

All authors conceived and wrote the paper; JEC and ADF conducted most analyses.

## ACKNOWLEDGEMENTS

We thank Marc Libault for critical reading of the manuscript and for inspiring this project. We are grateful to a number of colleagues who provided advice, answered questions, or provided unpublished materials: Jonathan Wendel, Andy Paterson, James Schnable, Pat Edger, Nicholas Panchy, and Shin-Han Shiu.

