## Supplemental Figures 1-4 for "Expression partitioning of duplicate genes at single cell resolution in *Arabidopsis* roots"

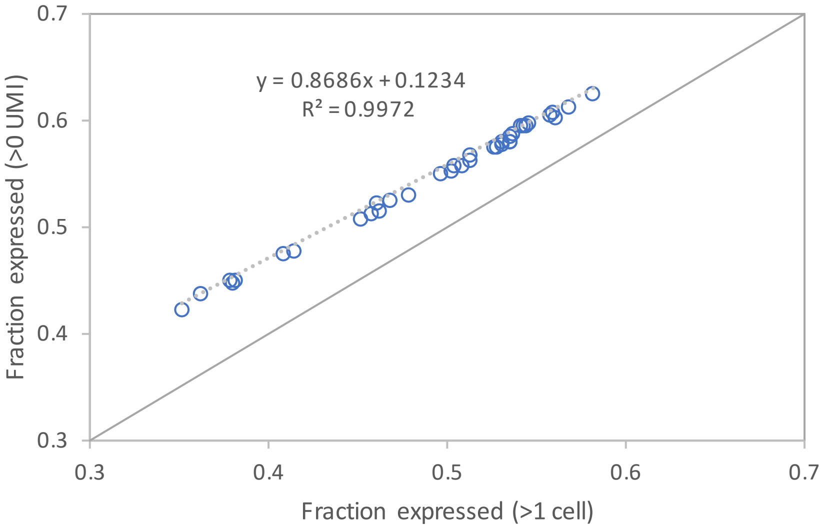


**Supplemental Figure 1.** Correlation between fractions of genes expressed per RCC calculated using two different thresholds of expression (≥1 UMI and >1 cell).

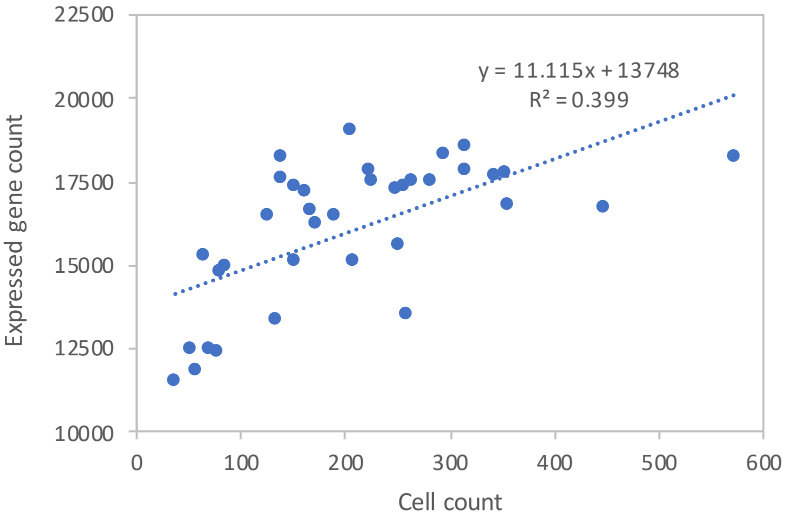


**Supplemental Figure 2.** Correlation between genes expressed per RCC and cells per RCC.

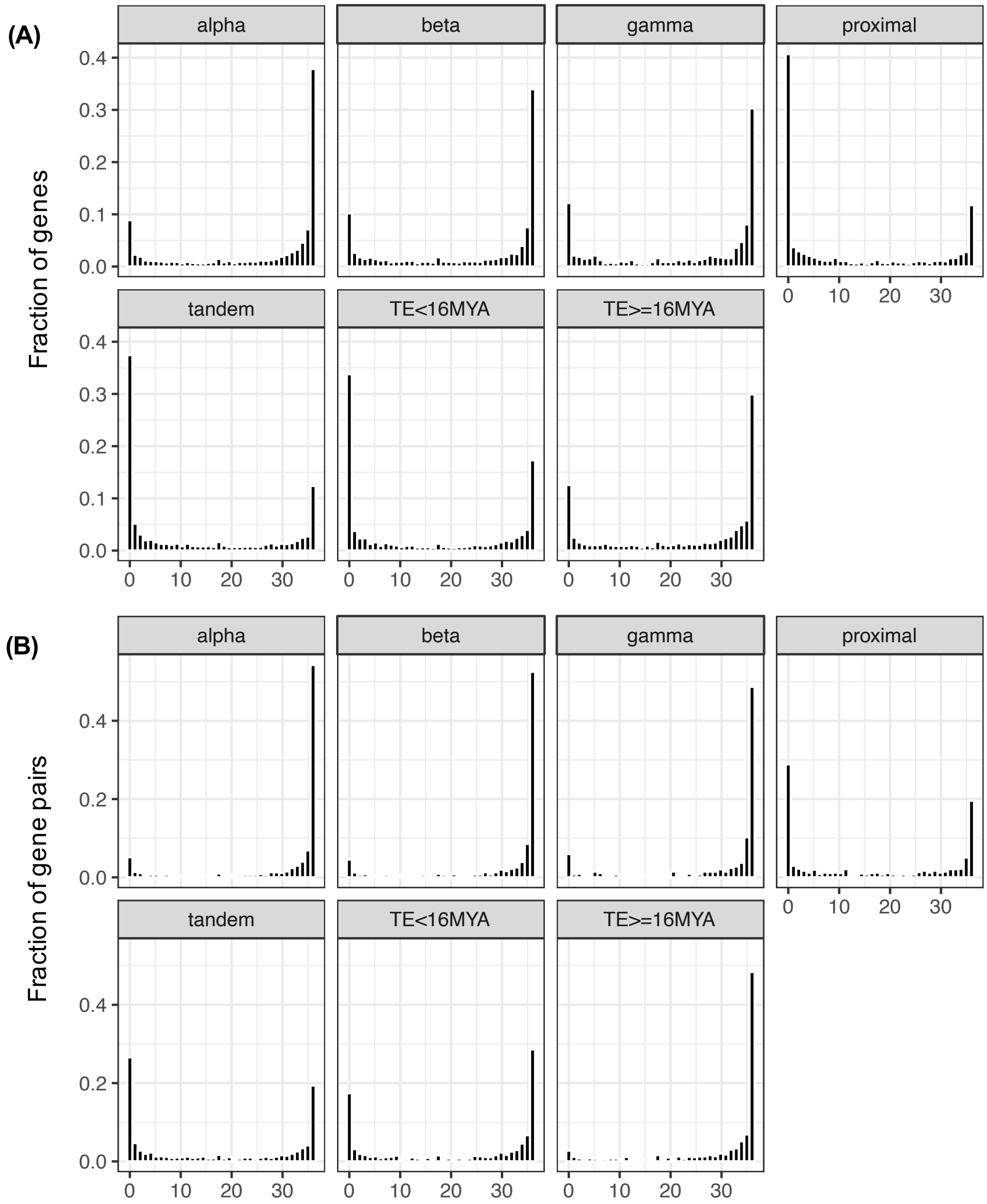


**Supplemental Figure 3.** Distributions of root cell clusters expressed per gene **(A**) or gene pair **(B)** for each duplication mechanism.

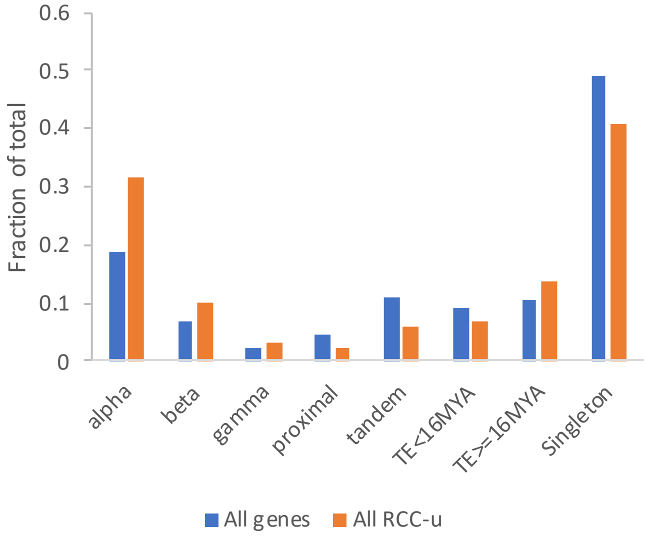


**Supplemental Figure 4.** RCC-u genes are enriched for WGD duplicates. Blue bars and orange bars indicate the fraction of genes that retain duplicates from the indicated mechanism for the whole genome and the RCC-u genes, respectively. The RCC-u gene set has a higher fraction of WGD duplicates and older TEs, and lower fraction of SSDs and singletons, relative to the whole genome.
