## Supplemental Tables 4-6, 8 for "Expression partitioning of duplicate genes at single cell resolution in *Arabidopsis* roots"

**Supplemental Table 4.** Counts of paralogous gene pairs by expression class and duplication type.

| Class | alpha | beta | gamma | proximal | tandem | TE<16MYA | TE≥16MYA | Total |
|---|---|---|---|---|---|---|---|---|
| 0 | 84 (2.7%) | 37 (2.6%) | 16 (3.2%) | 223 (28.8%) | 574 (27.1%) | 297 (17.5%) | 52 (2.8%) | 1,283 (11.2%) |
| 1 | 397 (12.8%) | 219 (15.4%) | 90 (17.8%) | 223 (28.8%) | 530 (25.0%) | 601 (35.5%) | 363 (19.5%) | 2,423 (21.1%) |
| 2 | 957 (30.9%) | 265 (18.7%) | 94 (18.6%) | 122 (15.8%) | 338 (15.9%) | 295 (17.4%) | 508 (27.3%) | 2,579 (22.5%) |
| 3 | 1570 (50.7%) | 814 (57.3%) | 283 (56.0%) | 192 (24.8%) | 641 (30.2%) | 473 (27.9%) | 860 (46.2%) | 4,833 (42.1%) |
| 4 | 88 (2.8%) | 85 (6.0%) | 22 (4.4%) | 14 (1.8%) | 38 (1.8%) | 27 (1.6%) | 78 (4.2%) | 352 (3.1%) |
| Total | 3096 | 1420 | 505 | 774 | 2121 | 1693 | 1861 | 11470 |

**Supplemental Table 5.** Counts of scB expression for Class I paralogues. A total of 2,423 pairs were assigned to Class 1 (scA expressed in > 1 cell/cluster in at least one cluster, scB expressed in ≤ 1 cell/cluster in all 36 RCCs).

| | | scB expressed at ≥5 reads in at least one SRA library | |
|---|---|---|---|
| scA expression cutoff[1] | Class 1 pairs | All SRA libraries | Excluding root libraries[2] |
| > 1 cell | 2,423 | 2,253 (93.0%) | 2,214 (91.4%) |
| > 1 cell, >1 RCC | 2,153 | 1,998 (92.8%) | 1,966 (91.3%) |
| > 1 cell, >5 RCC | 1,734 | 1,604 (92.5%) | 1,574 (90.8%) |

[1]Criteria for scA to be considered expressed in the root scRNA-seq libraries.
[2]121 out of 213 SRA libraries whose metadata indicate they were derived from root tissue or included root tissue (e.g., whole seedlings), were excluded from the analysis.

**Supplemental Table 6.** $K_a/K_s$ by expression Class and duplication type.

| Class | Duplication type | Count | Ka/Ks Mean | Ka/Ks SD |
|---|---|---|---|---|
| 0 | all | 1,283 | 0.79 | 1.06 |
| | alpha | 84 | 0.30 | 0.18 |
| | beta | 37 | 0.59 | 0.69 |
| | gamma | 16 | 0.46 | 0.26 |
| | proximal | 223 | 1.01 | 1.39 |
| | tandem | 574 | 0.75 | 0.61 |
| | TE<16 MYA | 297 | 0.96 | 1.58 |
| | TE≥16 MYA | 52 | 0.43 | 0.34 |
| 1 | all | 2,423 | 0.70 | 1.19 |
| | alpha | 397 | 0.39 | 0.45 |
| | beta | 219 | 0.44 | 0.64 |
| | gamma | 90 | 0.41 | 0.65 |
| | proximal | 223 | 0.87 | 0.96 |
| | tandem | 530 | 0.79 | 1.60 |
| | TE<16 MYA | 601 | 1.00 | 1.54 |
| | TE≥16 MYA | 363 | 0.52 | 0.52 |
| 2 | all | 2,577 | 0.44 | 0.95 |
| | alpha | 957 | 0.22 | 0.19 |
| | beta | 263 | 0.30 | 0.33 |
| | gamma | 94 | 0.45 | 0.61 |
| | proximal | 122 | 0.96 | 3.67 |
| | tandem | 338 | 0.52 | 0.56 |
| | TE<16 MYA | 295 | 0.66 | 0.82 |
| | TE≥16 MYA | 508 | 0.61 | 0.59 |
| 3 | all | 4,835 | 0.41 | 0.68 |
| | alpha | 1,570 | 0.27 | 0.32 |
| | beta | 816 | 0.31 | 0.47 |
| | gamma | 283 | 0.39 | 0.52 |
| | proximal | 192 | 0.65 | 0.82 |
| | tandem | 641 | 0.44 | 0.45 |
| | TE<16 MYA | 473 | 0.79 | 1.64 |
| | TE≥16 MYA | 860 | 0.50 | 0.49 |
| 4 | all | 352 | 0.30 | 0.23 |
| | alpha | 88 | 0.23 | 0.16 |
| | beta | 85 | 0.24 | 0.21 |
| | gamma | 22 | 0.31 | 0.29 |
| | proximal | 14 | 0.41 | 0.33 |
| | tandem | 38 | 0.33 | 0.22 |
| | TE<16 MYA | 27 | 0.29 | 0.15 |
| | TE≥16 MYA | 78 | 0.40 | 0.29 |

**Supplemental Table 8.** Breakdown by expression class and duplication mechanism of 4,599 Wang et al., (2013) paralogue pairs ubiquitously expressed in root cell clusters (RCC-u gene pairs).

| Duplication type | Class | Count | RCC-u mixed | RCC-u both |
|---|---|---|---|---|
| alpha | 0 | 0 | 0 | 0 |
| | 1 | 94 | 94 | 0 |
| | 2 | 590 | 148 | 442 |
| | 3 | 970 | 690 | 280 |
| | 4 | 31 | 17 | 14 |
| beta | 0 | 0 | 0 | 0 |
| | 1 | 60 | 60 | 0 |
| | 2 | 157 | 43 | 114 |
| | 3 | 494 | 371 | 123 |
| | 4 | 44 | 29 | 15 |
| gamma | 0 | 0 | 0 | 0 |
| | 1 | 24 | 24 | 0 |
| | 2 | 47 | 8 | 39 |
| | 3 | 165 | 131 | 34 |
| | 4 | 8 | 5 | 3 |
| proximal | 0 | 0 | 0 | 0 |
| | 1 | 34 | 34 | 0 |
| | 2 | 17 | 3 | 14 |
| | 3 | 95 | 83 | 12 |
| | 4 | 4 | 3 | 1 |
| tandem | 0 | 0 | 0 | 0 |
| | 1 | 50 | 50 | 0 |
| | 2 | 71 | 25 | 46 |
| | 3 | 279 | 233 | 46 |
| | 4 | 10 | 5 | 5 |
| TE<16MYA | 0 | 0 | 0 | 0 |
| | 1 | 140 | 140 | 0 |
| | 2 | 85 | 34 | 51 |
| | 3 | 249 | 212 | 37 |
| | 4 | 11 | 4 | 7 |
| TE≥16 MYA | 0 | 0 | 0 | 0 |
| | 1 | 103 | 103 | 0 |
| | 2 | 207 | 66 | 141 |
| | 3 | 528 | 406 | 122 |
| | 4 | 32 | 26 | 6 |
